# Gcn1 inhibits initiation of ribosome-associated quality control

**DOI:** 10.64898/2026.05.04.722640

**Authors:** Tristan Yew Kit Chan, Winifred Yun Xian Choo, Mi-Jeong Yoon, Young-Jun Choe

## Abstract

Ribosomes translating damaged mRNAs can stall, producing incomplete and potentially toxic polypeptides. The ribosome-associated quality control (RQC) pathway, triggered by collisions between stalled and trailing ribosomes, eliminates these aberrant translation products. Upon collision, ribosomes are split, and nascent polypeptides remaining on the large subunit are polyubiquitylated for proteasomal degradation. However, recent studies have shown that translation elongation rates vary along normal transcripts, leading to widespread ribosome collisions. Here, we investigated whether RQC is regulated to prevent unnecessary protein degradation during transient ribosome collisions. Using multiple conditions that induce transient ribosome pausing, including mild amino acid limitation, short polyadenosine tracts, and delayed translation termination in [*PSI*^+^] yeast cells, we show that the Gcn1-Gcn20 complex, a co-activator of the integrated stress response (ISR) kinase Gcn2, inhibits RQC initiation. We therefore propose that the Gcn1-Gcn20 complex has a dual role in ribosome collision responses: activating the ISR while inhibiting RQC. When this complex is impaired, physiological translation becomes more susceptible to RQC.

## Introduction

The α subunit of eukaryotic translation initiation factor 2 (eIF2α) is phosphorylated under various stress conditions (Costa-Mattioli & Walter, 2020; Pakos-Zebrucka *et al*, 2016; Ryoo, 2024). This modification reduces global protein synthesis while enabling selective translation of stress-responsive mRNAs, such as *GCN4* in yeast and *ATF4* in mammals (Dever *et al*, 2023; Hinnebusch, 2005; Wek, 2018). This conserved mechanism, known as the integrated stress response (ISR), promotes cellular adaptation and survival under stress. In mammals, the ISR is triggered by multiple eIF2α kinases, each activated by specific stress signals (Fessler *et al*, 2020; Guo *et al*, 2020; Harding *et al*, 1999). In contrast, yeast relies on a single known ISR kinase, Gcn2, which can be activated by amino acid starvation (Dever *et al*, 1992; Wek *et al*, 1989). When amino acids are limited, uncharged tRNAs build up and bind Gcn2 through its histidyl-tRNA synthetase-like domain (Dong *et al*, 2000; Wek *et al*, 1995; Zaborske *et al*, 2009). Although the domain lost aminoacylation activity, structural elements to bind the acceptor stem and anticodon loop of uncharged tRNAs are maintained (Bou-Nader *et al*, 2024). In addition to uncharged tRNAs, the ribosomal P-stalk has also been identified as a Gcn2-activating ligand (Harding *et al*, 2019; Inglis *et al*, 2019), suggesting a role of the ribosome in Gcn2 activation. Indeed, the ribosome-associated Gcn1-Gcn20 complex has long been known as a critical Gcn2 co-activator (Garcia-Barrio *et al*, 2000; Marton *et al*, 1997). Recent studies have identified how translating ribosomes set the stage to activate Gcn2. Failed tRNA charging due to amino acid starvation results in stalled ribosomes, which can collide with trailing ribosomes (Darnell *et al*, 2018; Meydan & Guydosh, 2020; Wu *et al*, 2020). Then the Gcn1-Gcn20 complex associates with these collided ribosomes and recruits Gcn2 for its activation (Paternoga *et al*, 2025; Pochopien *et al*, 2021; Wu *et al*., 2020; Yan & Zaher, 2021; Zhou *et al*, 2025). Consistently, amino acid starvation is not the only condition that activates Gcn2. Other stressors that induce ribosome stalling, such as translation inhibitors, UV irradiation, or mRNA alkylation, can also activate Gcn2 (Ishimura *et al*, 2016; Wu *et al*., 2020; Yan & Zaher, 2021). Although the precise mechanism of Gcn2 activation remains to be fully elucidated (Gupta & Hinnebusch, 2023; Misra *et al*, 2024; Paternoga *et al*., 2025), the Gcn1-Gcn20 complex plays a central role in this process by binding collided di-ribosomes (or disomes).

Ribosomal stalling generates incomplete polypeptides that can be toxic to cells. To prevent their accumulation, eukaryotes employ a specialized co-translational protein degradation pathway, ribosome-associated quality control (RQC) (Brandman *et al*, 2012; Inada, 2026; Joazeiro, 2019; Sitron & Brandman, 2020). When ribosomes stall and collide, the E3 ligase Hel2 in yeast (or ZNF598 in mammals) ubiquitylates the small subunits of disomes (Garzia *et al*, 2017; Ikeuchi *et al*, 2019; Juszkiewicz *et al*, 2018; Juszkiewicz & Hegde, 2017; Matsuo *et al*, 2017; Simms *et al*, 2017; Sundaramoorthy *et al*, 2017). This ubiquitin serves as a signal to recruit the RQC-trigger (RQT) complex in yeast (or the ASC-1 complex in mammals), which resolves the collided disomes (Juszkiewicz *et al*, 2020; Matsuo *et al*, 2020). Unlike canonical translation termination and ribosome recycling at stop codons, ribosome splitting by the RQT complex does not involve peptidyl-tRNA hydrolysis (Best *et al*, 2023). As a result, the dissociated large ribosomal subunits remain bound to a peptidyl-tRNA, and this aberrant molecular complex is recognized by Rqc2 (Fig. EV1A) (Lyumkis *et al*, 2014; Shao *et al*, 2015; Shen *et al*, 2015). Rqc2 then recruits the E3 ligase Ltn1, which polyubiquitylates the stalled polypeptides, targeting them for proteasomal degradation (Bengtson & Joazeiro, 2010; Doamekpor *et al*, 2016; Shao *et al*., 2015; Shen *et al*., 2015).

RQC is highly sensitive in detecting disomes within vast translation pools, raising the question of whether and how transiently collided ribosomes—resulting from elongation rate heterogeneity along transcripts (Aguilar Rangel *et al*, 2024; Collart & Weiss, 2020; Hanson & Coller, 2018; Ingolia *et al*, 2019; Komar *et al*, 2024)—avoid the degradation of otherwise normal nascent polypeptides. In this study, we investigate conditions that induce transient ribosome pausing rather than persistent stalling and show that the Gcn1-Gcn20 complex inhibits Hel2-dependent RQC initiation, thereby fine-tuning co-translational protein degradation.

## Results

### Amino acid starvation induces uS10 ubiquitylation

Amino acid starvation impairs tRNA aminoacylation (Darnell *et al*., 2018; Zaborske *et al*., 2009), causing ribosome stalling and collisions that activate the ISR (Wu *et al*., 2020; Yan & Zaher, 2021). Gcn1 is a collision sensor (Fig. EV1B) that activates the ISR kinase Gcn2 on the ribosome (Pochopien *et al*., 2021; Wu *et al*., 2020; Yan & Zaher, 2021). We asked whether ribosome-associated ISR factors also influence RQC. As a first step, we examined whether the ISR-inducing starvation condition can also trigger RQC. RQC initiation upon ribosome stalling requires Hel2-mediated ubiquitylation of the small subunit protein uS10 (Fig. EV1C) (Garzia *et al*., 2017; Juszkiewicz & Hegde, 2017; Matsuo *et al*., 2017; Sundaramoorthy *et al*., 2017). Although structural information on Hel2 recognition of collided ribosomes remains lacking, Hel2 preferentially targets collided ribosomes rather than isolated stalled monosomes (Ikeuchi *et al*., 2019; Juszkiewicz *et al*., 2018). Consistent with this collision dependence, uS10 ubiquitylation was more efficient at lower cycloheximide concentrations (∼1 μg/ml) than at higher concentrations (Fig. EV1D), in line with previous studies using other ribosome inhibitors in mammalian cells (Juszkiewicz *et al*., 2018). Although multiple proteins of the small ribosome subunit can be ubiquitylated, uS10 ubiquitylation is the only modification known to be essential for RQC initiation in yeast (Matsuo *et al*., 2017). As shown previously, mutation of the uS10 ubiquitylation sites disrupted RQC (Fig. EV1E,F) (Matsuo *et al*., 2017). CGA codon repeats induce strong ribosome stalling in yeast (Letzring *et al*, 2010), and the resulting stalled polypeptides can be detected in RQC-defective *ltn1*Δ cells (Fig. EV1F, lane 1). However, when uS10 ubiquitylation was prevented, RQC was not initiated, allowing ribosomes to read through the CGA codon repeat (Fig. EV1F, lane 2).

We next investigated whether amino acid starvation can induce uS10 ubiquitylation. Although previous studies showed that ISR-inducing translation stress can induce uS3 ubiquitylation (Meydan & Guydosh, 2020; Yan & Zaher, 2021), this modification does not necessarily indicate RQC activation. First, uS3 ubiquitylation is mediated primarily by the E3 ligase Mag2 rather than Hel2 (Sugiyama *et al*, 2019). uS3 ubiquitylation was still detected in *hel2*Δ cells but was abolished in *mag2*Δ cells (Fig. EV1G, lanes 4 and 7). Even when *HEL2* was overexpressed, uS3 ubiquitylation was not induced in the absence of Mag2 (Fig. EV1G, lane 8). Conversely, *MAG2* overexpression enhanced uS3 ubiquitylation whether Hel2 was present or absent (Fig. EV1G, lanes 3 and 6). Second, uS3 ubiquitylation is dispensable for RQC. Even when uS3 ubiquitylation was abolished by mutation of its modification site (Fig. EV1H) (Matsuo *et al*., 2017), RQC was initiated normally, as shown by generation of the stalled protein (Fig. EV1I, lane 4), in stark contrast to the RQC impairment caused by loss of uS10 ubiquitylation (Fig. EV1I, lane 2). Accordingly, we revisited whether amino acid starvation activates RQC by assessing uS10 ubiquitylation. Because the yeast strains used in this study were leucine auxotrophs, we depleted leucine from the culture media. Leucine depletion effectively activated the ISR, as shown by increased eIF2α phosphorylation, but did not induce uS10 ubiquitylation (Fig. 1A, lanes 1 and 3). Thus, it remained unclear whether starvation-induced ribosome stalling is targeted by RQC. Previous studies have shown that ISR activation can antagonize RQC (Yan & Zaher, 2021), likely because ISR-mediated repression of global translation reduces ribosome loading and thereby lowers the frequency of ribosome collisions. To test this possibility, we disrupted the ISR by mutating the phosphorylation site in eIF2α to alanine (S52A) and reassessed uS10 ubiquitylation. Indeed, this ISR-defective strain exhibited increased uS10 ubiquitylation upon leucine depletion (Fig. 1A, lanes 2 and 4), indicating that starvation-induced stalled ribosomes can be recognized by the RQC pathway.

**Figure 1.**
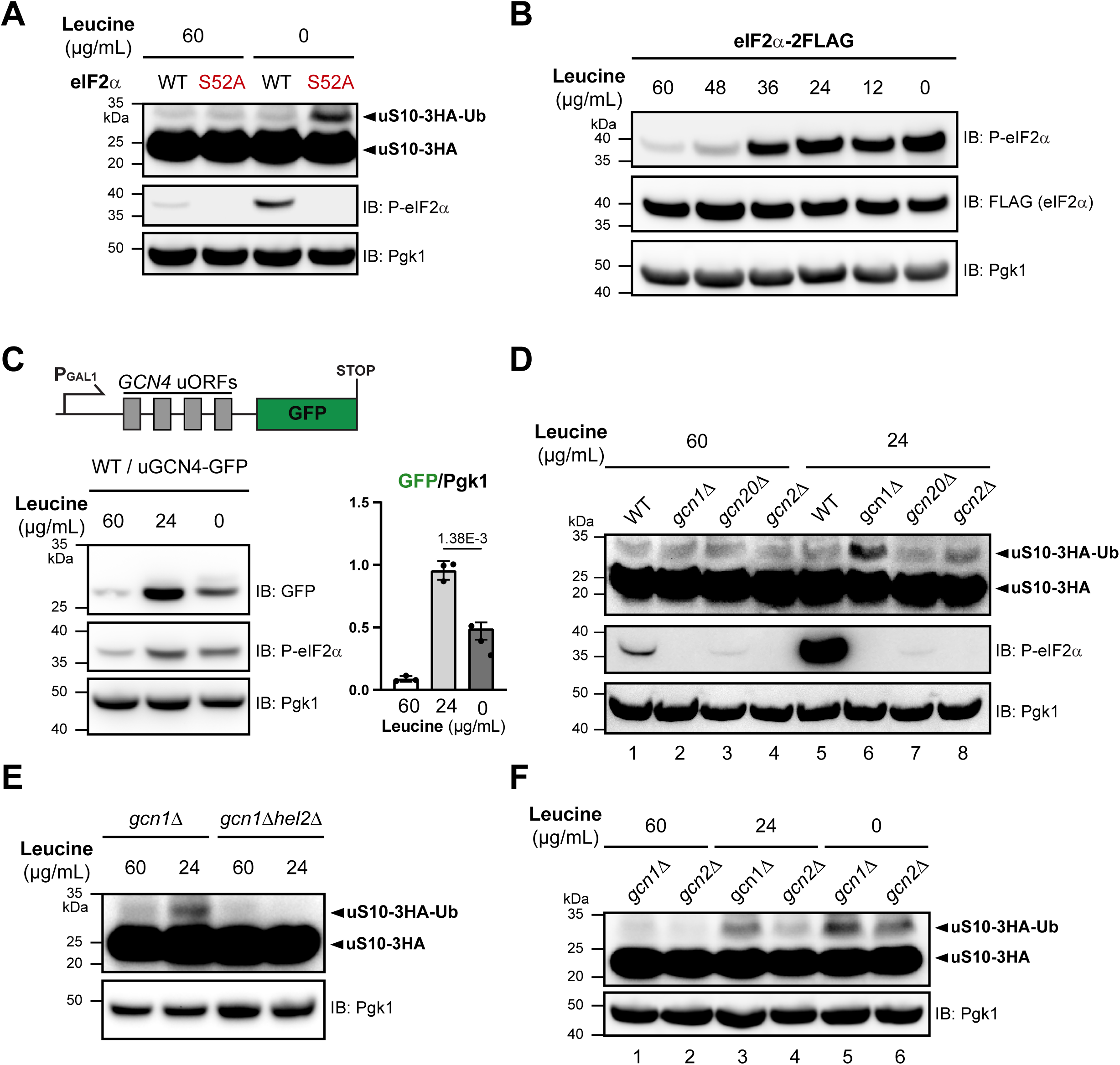
Gcn1 inhibits RQC during partial amino acid depletion (A) Genomic uS10/*RPS20* was C-terminally tagged with a 3×HA epitope. To impair the ISR, the S52A mutation was introduced into genomic eIF2α/*SUI2*. Cells precultured in nutrient-rich medium were collected by centrifugation, resuspended in complete or leucine-deficient medium, and incubated for 15 min. uS10 ubiquitylation and ISR activation were assessed by immunoblotting with antibodies against HA and phosphorylated eIF2α (P-eIF2α), respectively. Pgk1 served as a loading control. IB, immunoblotting; Ub, ubiquitin. (B) The genomic eIF2α/*SUI2* locus was C-terminally tagged with a 2×FLAG epitope. Cells precultured in nutrient-rich medium were collected by centrifugation, resuspended in media containing the indicated leucine concentrations, and incubated for 15 min. ISR activation was assessed by immunoblotting with an antibody against phosphorylated eIF2α (P-eIF2α). (C) The ISR reporter, containing the four *GCN4* uORFs upstream of the GFP start codon, was expressed in galactose media with the indicated leucine concentrations for 3 h. GFP levels were analyzed by immunoblotting and normalized to Pgk1. GFP/Pgk1 ratios are shown in the bar graph. The *P* value was calculated using a paired, two-tailed t-test. Black dots, individual data points (*n* = 3); error bars, SEM. (D-F) *GCN* genes or *HEL2* were deleted in the uS10-3HA strain. Cells precultured in nutrient-rich medium were collected by centrifugation, resuspended in media containing the indicated leucine concentrations, and incubated for 15 min. uS10 ubiquitylation was assessed by anti-HA immunoblotting.

### Gcn1 inhibits Hel2-dependent uS10 ubiquitylation

We next asked whether mild amino acid limitation, rather than complete starvation, can provoke cellular responses. We therefore titrated leucine in the culture media and found that partial leucine depletion was sufficient to trigger eIF2α phosphorylation (Fig. 1B). The cellular response to partial leucine limitation (24 μg/ml) was rapid, becoming evident within 5 min of stress (Fig. EV1J). However, overall translation remained relatively active at this leucine concentration, as indicated by the preservation of robust polysomes (Fig. EV1K; compare 24 μg/ml and 0 μg/ml leucine). To validate the functional consequence of eIF2α phosphorylation under mild starvation (Fig. 1B, 24 μg/ml leucine), we examined translational control mediated by the four upstream open reading frames (uORFs) of *GCN4* (Dever *et al*., 2023; Hinnebusch, 2005; Hinnebusch *et al*, 2016). *GCN4* encodes the key ISR transcription factor that activates amino acid biosynthetic genes. After translation of the first uORF, the 40S subunit can remain bound to the mRNA and resume scanning for reinitiation. Under normal conditions, the 40S subunit reinitiates translation at downstream uORFs, and ribosomes dissociate from the mRNA at their stop codons, preventing translation of the main *GCN4* ORF. Upon ISR activation, eIF2α phosphorylation reduces reinitiation at downstream uORFs, allowing the scanning 40S subunit to bypass them and initiate translation at the main *GCN4* ORF. To determine whether the observed eIF2α phosphorylation under partial leucine limitation (Fig. 1B) activates *GCN4*-type translational control, we designed a GFP reporter whose translation is regulated by the *GCN4* uORFs (Fig. 1C). As expected, GFP was barely detectable in leucine-rich media (Fig. 1C, lane 1), confirming that the *GCN4* uORFs suppress downstream GFP translation. In contrast, GFP translation was activated by leucine limitation (Fig. 1C, lanes 2 and 3), demonstrating that the observed eIF2α phosphorylation engaged *GCN4*-type translational control. Notably, GFP expression was significantly higher under partial leucine depletion than under complete leucine starvation (Fig. 1C, compare lanes 2 and 3), consistent with the greater overall translational activity observed at 24 μg/ml leucine than at 0 μg/ml leucine (Fig. EV1K).

Together, these data indicate that leucine limitation at 24 μg/ml induces the ISR (Fig. 1B,C) without severely disrupting polysomes (Fig. EV1K). As expected, eIF2α phosphorylation under this condition required the canonical yeast ISR factors Gcn1, Gcn20, and Gcn2 (Fig. 1D, compare lane 5 with lanes 6–8). We then examined RQC activity in cells lacking these ISR factors, anticipating enhanced uS10 ubiquitylation, as observed in the eIF2α phosphorylation site mutant (Fig. 1A, lane 4). Strikingly, however, only *gcn1*Δ cells exhibited increased uS10 ubiquitylation under this mild stress condition (Fig. 1D, lane 6). In contrast, deletion of the other ISR factors, *GCN20* or *GCN2*, did not noticeably increase uS10 ubiquitylation (Fig. 1D, lanes 7 and 8). Thus, the enhanced uS10 ubiquitylation in *gcn1*Δ cells was not simply attributable to the ISR impairment. In the absence of Hel2, uS10 ubiquitylation was abolished (Fig. 1E), indicating that the enhanced uS10 ubiquitylation in *gcn1*Δ cells was mediated by Hel2, not by other E3 ligases. Moreover, cycloheximide at a high concentration effectively suppressed uS10 ubiquitylation in low-leucine media (Fig. EV1L), consistent with the preferential targeting of collided ribosomes, rather than isolated stalled monosomes, by Hel2. Notably, Hel2 abundance did not increase in *gcn1*Δ cells (Fig. EV1M), suggesting that the enhanced uS10 ubiquitylation was not caused by increased Hel2 levels. Instead, collided ribosomes in *gcn1*Δ cells may be more readily engaged by Hel2 and/or more efficiently ubiquitylated on uS10. When leucine was completely depleted, however, both *gcn1*Δ and *gcn2*Δ cells displayed similarly robust uS10 ubiquitylation (Fig. 1F, lanes 5 and 6). Thus, Gcn1 effectively inhibits Hel2-dependent uS10 ubiquitylation under mild amino acid limitation, when translation remains sufficiently active.

### The Gcn1-Gcn20 complex inhibits RQC during translation of polyadenosine tracts

We next asked whether the increased Hel2-dependent uS10 ubiquitylation observed in *gcn1*Δ cells leads to enhanced RQC. In addition, it remained unclear whether Gcn1 regulates Hel2 activity only during amino acid depletion, as the uS10 ubiquitylation assay may not be sensitive enough to detect subtle differences under leucine-replete conditions. To address these questions, we used polyadenosine tracts, which induce ribosome pausing independently of amino acid supply. Polyadenosine tracts form a helical structure within translating ribosomes, thereby hindering decoding (Chandrasekaran *et al*, 2019; Tesina *et al*, 2020). This translation-inhibitory effect increases with tract length (Arthur *et al*, 2015), providing a means to tune stalling strength. We compared two polyadenosine tract lengths inserted between GFP and mCherry (Fig. 2A). As expected, a tract of ten consecutive AAA codons (30 nucleotides) generated more stalled GFP than a tract of eight AAA codons (24 nucleotides) in *ltn1*Δ cells (Fig. 2A, lanes 2 and 4). In contrast, stalled GFP was barely detectable in wild-type (WT) cells, indicating that these stalled products were degraded through the Ltn1-mediated RQC pathway.

**Figure 2.**
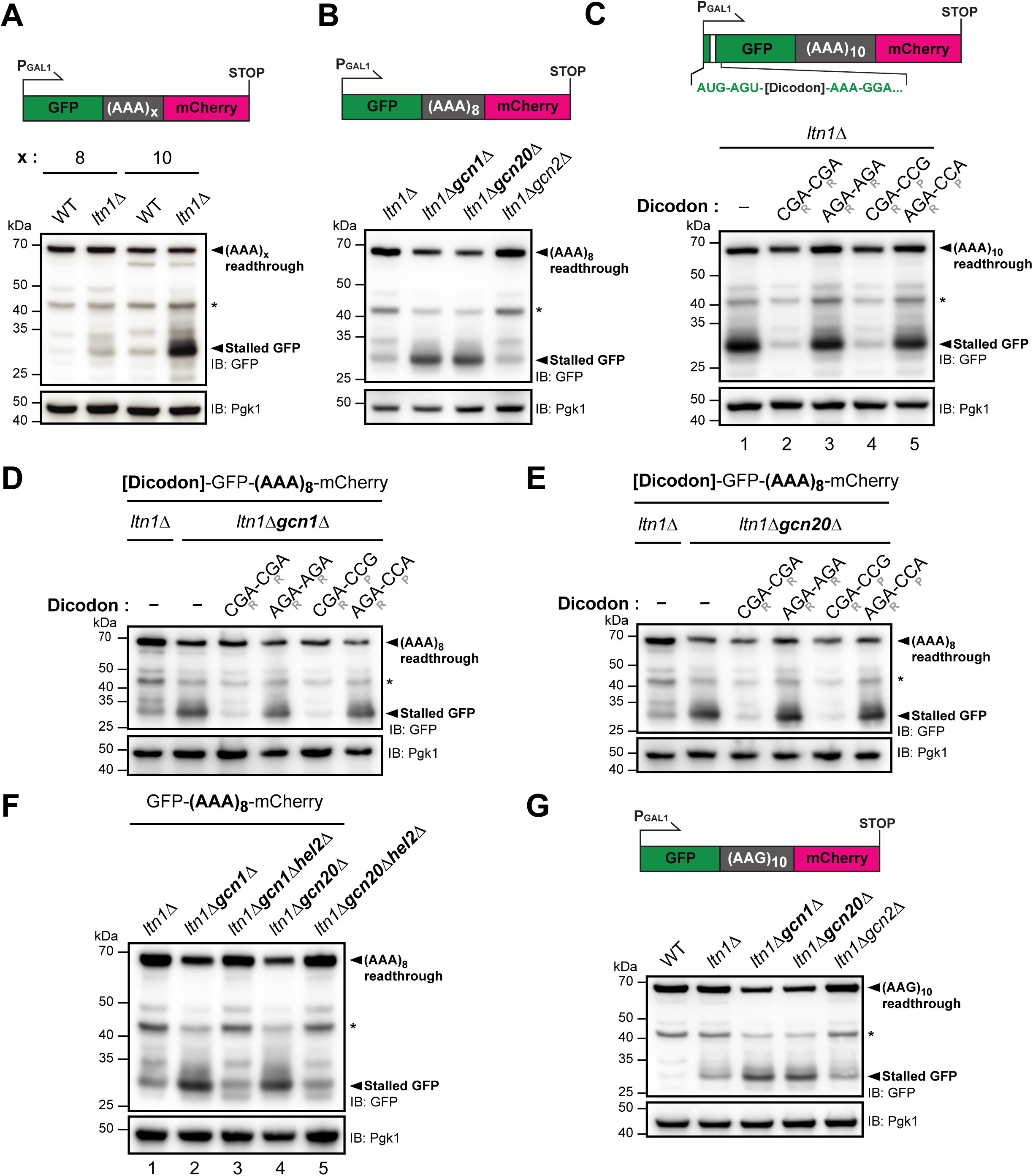
The Gcn1-Gcn20 complex inhibits RQC during translation of polyadenosine tracts (A) In the polyadenosine reporter schematic, (AAA)_x_ denotes x repeats of the AAA codon. Reporters were expressed from the *GAL1* promoter in WT or *ltn1*Δ cells. Cell lysates were analyzed by anti-GFP immunoblotting. Pgk1 served as a loading control. IB, immunoblotting; * degradation product of the readthrough protein. (B) The (AAA)_8_ reporter, corresponding to x = 8 in the schematic in (A), was expressed in the indicated strains. Cell lysates were analyzed by anti-GFP immunoblotting. (C) Translation-inhibitory dicodons (CGA-CGA and CGA-CCG) or their synonymous counterparts (AGA-AGA and AGA-CCA) were inserted at the third codon position of the (AAA)_10_ reporter (x = 10 in (A)), following AUG (start codon) and AGU. CGA and AGA encode arginine (R); CCG and CCA encode proline (P). Reporters were expressed in *ltn1*Δ cells. (D-E) The translation-inhibitory dicodons and their synonymous controls were inserted into the (AAA)_8_ reporter shown in (B), at the same position described in (C). Reporters were expressed in *ltn1*Δ*gcn1*Δ (D) and *ltn1*Δ*gcn20*Δ (E) cells. The (AAA)_8_ reporter lacking dicodon insertion was analyzed in lanes 1 and 2 of both panels. (F-G) The (AAA)_8_ reporter (F) and (AAG)_10_ reporter (G) were expressed under the *GAL1* promoter in the indicated strains. Cell lysates were analyzed by anti-GFP immunoblotting.

We next asked whether Gcn1 influences RQC. Accordingly, we expressed the weak (AAA)_8_ reporter in cells lacking individual ISR factors: *GCN1*, *GCN20*, or *GCN2*. Remarkably, stalled GFP levels increased substantially in *gcn1*Δ and *gcn20*Δ cells, concomitant with reduced (AAA)_8_ readthrough (Fig. 2B, lanes 2 and 3). However, stalled GFP was not detected in cells expressing Ltn1, even when *GCN1* or *GCN20* was deleted (Fig. EV2A, compare lanes 2 and 3, and lanes 4 and 5). In contrast to the reduced (AAA)_8_ readthrough in *gcn1*Δ and *gcn20*Δ cells (Fig. 2B), a control reporter lacking the polyadenosine tract produced comparable levels of full-length protein in these mutant cells and did not generate stalled GFP (Fig. EV2B, lanes 2 and 3). Together, these results suggest that, when *GCN1* or *GCN20* is deleted, ribosomes translating the short (AAA)_8_ tract undergo RQC-associated splitting, and the resulting GFP product is processed by the 60S-associated E3 ligase Ltn1. Importantly, all experiments using polyadenosine reporters in this study were performed in amino acid-rich media. Thus, the enhanced RQC observed in *gcn1*Δ and *gcn20*Δ cells is unlikely to result from the ISR impairment. Consistently, deletion of *GCN2*, which encodes the only ISR kinase in yeast, did not affect RQC of the (AAA)_8_ reporter (Fig. 2B, lane 4).

Collectively, the increased uS10 ubiquitylation observed in *gcn1*Δ cells under amino acid limitation (Fig. 1D,F) is consistent with their enhanced RQC during polyadenosine translation. In contrast, increased uS10 ubiquitylation was not detected in *gcn20*Δ cells (Fig. 1D, lane 7), despite their strong RQC phenotype with polyadenosine reporters (Fig. 2B, lane 3). Previous studies have shown that Gcn20 is not absolutely required for ISR activation (Vazquez de Aldana *et al*, 1995), consistent with the ability of Gcn1 to bind ribosomes independently of Gcn20 (Marton *et al*., 1997). Thus, in *gcn20*Δ cells, Gcn1 alone may still partially inhibit Hel2-dependent uS10 ubiquitylation under amino acid stress conditions. However, efficient recognition of rare, transient ribosome collisions during translation of short polyadenosine reporters may require the complete Gcn1-Gcn20 complex.

### Enhanced RQC upon loss of the Gcn1-Gcn20 complex requires ribosome collision

Although the Gcn1-Gcn20 complex recognizes collided ribosomes, sucrose gradient analyses detected both Gcn1 and Gcn20 not only in disome fractions but also in monosome fractions (Marton *et al*., 1997; Sattlegger & Hinnebusch, 2000, 2005; Yan & Zaher, 2021; Zhou *et al*., 2025). Furthermore, Gcn1-selective ribosome profiling enriched ribosome-protected mRNA fragments of both monosome and disome sizes (Muller *et al*, 2023). We therefore asked whether the Gcn1-Gcn20 complex could function on stalled monosomes to inhibit potential collision-independent RQC initiation. To address this question, we sought to reduce ribosome loading onto our reporters, thereby lowering the probability of ribosome collision and enriching for stalled ribosomes in the monosome state. Notably, certain codon-anticodon interactions in the ribosomal P-site are known to influence decoding at the A-site (Gamble *et al*, 2016). CGA-CGA and CGA-CCG are such inhibitory dicodons: the CGA codon-anticodon interaction in the P-site slows decoding of the following CGA or CCG codon in the A-site. We inserted these inhibitory dicodons near the start codon to slow ribosomes shortly after initiation (Fig. 2C, schematic). This start-proximal delay was expected both to occlude the initiation region, reducing further ribosome loading, and to increase the spacing between the dicodon-delayed ribosome and downstream elongating ribosomes, thereby lowering the probability of collision. We first verified the translation-inhibitory effect of the CGA-CGA and CGA-CCG dicodons by inserting them into a non-stalling reporter. Indeed, these dicodons reduced reporter expression (Fig. EV2C, lanes 2 and 4), whereas the synonymous dicodons AGA-AGA and AGA-CCA did not (Fig. EV2C, lanes 3 and 5). Thus, the reduced reporter levels were attributable to impaired translation rather than destabilization caused by the encoded dipeptide insertions, Arg-Arg and Arg-Pro. The translation-inhibitory dicodons and their synonymous controls were then introduced into the (AAA)_10_ reporter (Fig. 2C). Remarkably, insertion of just two codons, either CGA-CGA or CGA-CCG, near the start codon strongly reduced RQC initiation at the (AAA)_10_ tract located ∼240 codons downstream, as shown by the markedly reduced stalled GFP signal (Fig. 2C, lanes 2 and 4). This weak signal was not simply due to reduced reporter translation. Even when lysates from cells expressing the inhibitory-dicodon reporters were loaded at threefold higher amounts (Fig. EV2D, lanes 4 and 5), stalled GFP levels remained much lower than in the control reporter lacking these dicodons (Fig. EV2D, lane 1).

We next examined whether *gcn1*Δ and *gcn20*Δ cells could efficiently initiate RQC even when stalled ribosomes were biased toward the monosome state by the inhibitory dicodons. We modified the (AAA)_8_ reporter in Fig. 2B by introducing the dicodons and expressed them in *gcn1*Δ (Fig. 2D) and *gcn20*Δ cells (Fig. 2E). Insertion of the inhibitory dicodons CGA-CGA or CGA-CCG prevented accumulation of stalled GFP, unlike the synonymous dicodon controls AGA-AGA and AGA-CCA. The weak stalled GFP signals from the inhibitory-dicodon reporters were not simply due to reduced overall reporter translation. Even when higher amounts of lysates from cells expressing these reporters were loaded, stalled GFP levels remained substantially lower than those from the no-dicodon control reporter (Fig. EV2E). Thus, when ribosome loading was reduced and stalled ribosomes were biased toward the monosome state, RQC remained inefficient even in *gcn1*Δ and *gcn20*Δ cells. This argues against broad RQC recognition of stalled monosomes upon loss of the Gcn1-Gcn20 complex and instead supports a model in which these mutants are more sensitive to ribosome collisions. Consistently, deletion of the collision sensor *HEL2* suppressed RQC in *gcn1*Δ and *gcn20*Δ cells (Fig. 2F; compare lanes 2 and 3, and lanes 4 and 5).

We next asked whether the enhanced RQC observed in *gcn1*Δ and *gcn20*Δ cells is restricted to polyadenosine-mediated stalling or extends to other forms of ribosome stalling. To inhibit elongation by a distinct mechanism, we used reporters encoding polylysine tracts, whose positive charge inhibits translation through interactions with the negatively charged rRNA backbone in the ribosome exit tunnel (Dimitrova *et al*, 2009; Lu & Deutsch, 2008). A 10-residue polylysine tract encoded by AAG codons induced mild ribosome pausing, generating stalled GFP that accumulated to higher levels in cells lacking the Gcn1-Gcn20 complex (Fig. 2G). However, deletion of the Gcn2 kinase or the recently identified ISR factor Mbf1 did not enhance stalled GFP accumulation (Figs. 2G and EV2F–H) (Kim *et al*, 2024). In summary, our data suggest that the Gcn1-Gcn20 complex limits Hel2-dependent RQC initiation at collided ribosomes, especially when collisions arise from transient ribosome pausing.

### Compromised translation termination in [*PSI*^+^] cells induces ribosome collisions and activates the ISR

Having examined mild ribosome pausing during elongation, induced by partial leucine limitation and short polyadenosine tracts, we next asked whether the Gcn1-Gcn20 complex also limits RQC during a distinct form of pausing caused by inefficient translation termination. To this end, we used the yeast prion [*PSI*^+^] (Chernoff *et al*, 1995; Patino *et al*, 1996). In [*PSI*^+^] cells, the eukaryotic release factor 3 (eRF3), encoded by *SUP35*, forms self-propagating amyloid-like aggregates (Glover *et al*, 1997; Tanaka *et al*, 2004). This aggregation reduces the pool of functional eRF3, thereby delaying translation termination at stop codons. This delay allows near-cognate tRNAs to decode stop codons, thereby promoting stop codon readthrough (Blanchet *et al*, 2014). Stop codon readthrough in [*PSI*^+^] cells has been widely monitored using the *ade1-14* allele, which carries a nonsense mutation (Fig. EV3A). *ADE1* encodes an enzyme in the adenine biosynthesis pathway. In [*psi*⁻] cells, soluble Sup35 efficiently terminates translation at the premature stop codon in *ade1-14*, yielding a truncated, nonfunctional Ade1 product. Consequently, these cells fail to grow on media lacking adenine (Fig. 3A, top row of the -Adenine plate). When adenine is supplied at limiting levels, the cells can grow but accumulate intermediates in the adenine biosynthesis pathway, leading to red pigmentation (Fig. 3A, ¼YPD) (Smirnov *et al*, 1967). In [*PSI*^+^] cells, readthrough of the *ade1-14* nonsense mutation allows production of functional Ade1, thereby supporting growth on adenine-deficient media and preventing red pigmentation (Fig. 3A, second row of the spotting assay). Although stop codon readthrough has been extensively studied in [*PSI*^+^] cells, whether Sup35 aggregation also promotes ribosome collisions at stop codons remains unexplored.

**Figure 3.**
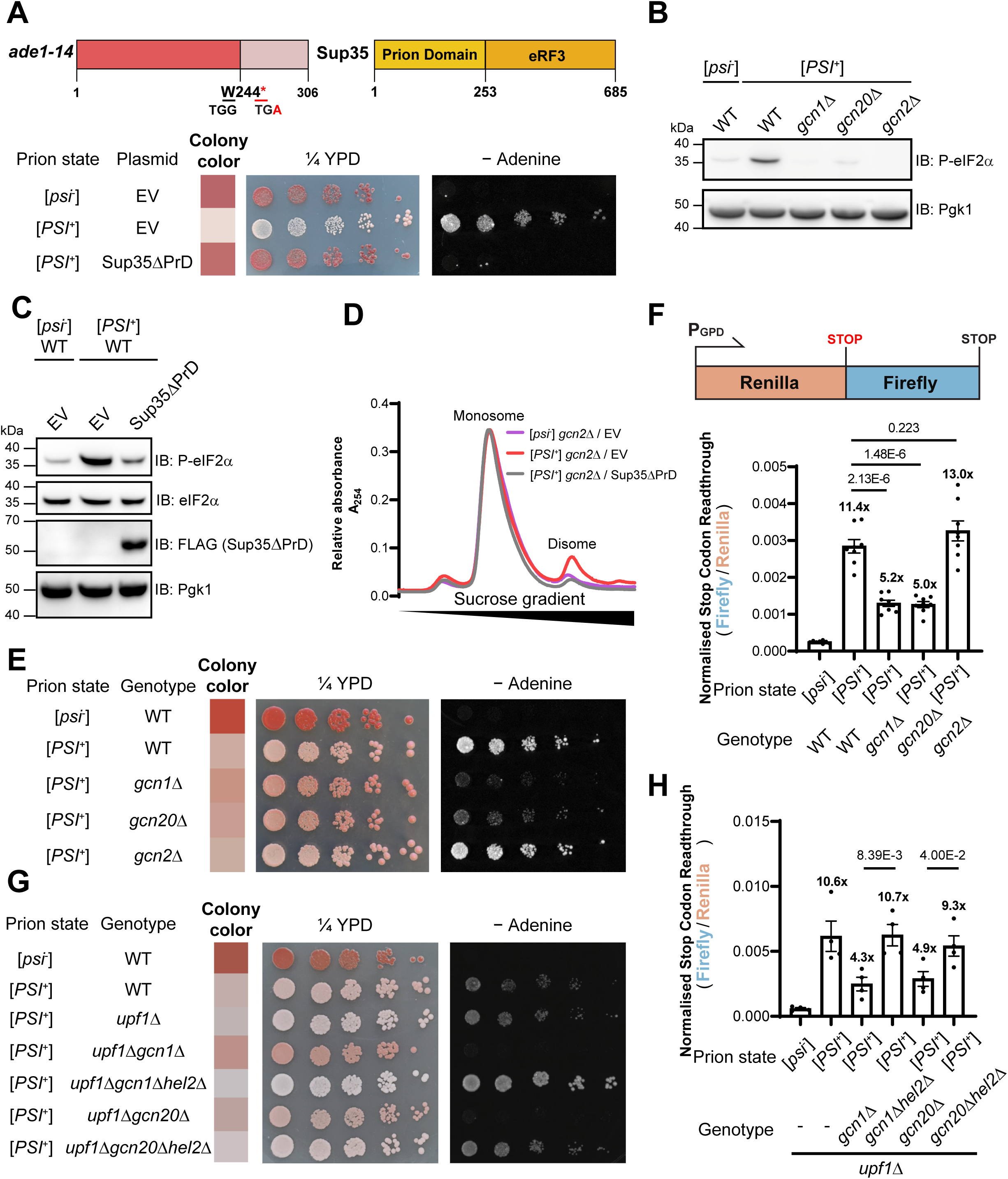
The Gcn1-Gcn20 complex preserves stop codon readthrough in [*PSI*^+^] cells (A) The nonsense mutation in *ade1-14* and the domain organization of Sup35 are shown. The *ade1-14* allele carries a TGG-to-TGA nonsense mutation at codon 244 of *ADE1*. Sup35 lacking the N-terminal prion domain (Sup35ΔPrD) was C-terminally FLAG-tagged and expressed from a plasmid under the native *SUP35* promoter. Empty vector (EV) served as a control. Cells carrying the plasmids were cultured in vector-selective media, fivefold serially diluted, and spotted onto adenine-limited (¼YPD) and adenine-deficient (−Adenine) plates. ¼YPD contains one-quarter of the yeast extract present in standard YPD medium and therefore imposes stronger adenine limitation. (B) Cells of the indicated genotypes and [*psi*⁻]/[*PSI*⁺] states were cultured in nutrient-rich synthetic complete media under normal conditions. Cell lysates were analyzed by immunoblotting (IB) using an antibody against phosphorylated eIF2α (P-eIF2α). Pgk1 served as a loading control. (C) [*psi*⁻] and [*PSI*⁺] cells transformed with empty vector or a Sup35ΔPrD expression vector were cultured in nutrient-rich vector-selective media under normal conditions. Cell lysates were analyzed by immunoblotting for P-eIF2α. As in (A), Sup35ΔPrD was C-terminally FLAG-tagged and expressed from the native *SUP35* promoter. (D) [*psi*⁻] *gcn2*Δ and [*PSI*⁺] *gcn2*Δ cells transformed with empty vector or a Sup35ΔPrD expression vector were cultured in nutrient-rich vector-selective media under normal conditions. Cell lysates were treated with P1 nuclease. Normalized lysates were layered onto 7–47% sucrose gradients, and monosome/disome profiles were monitored by absorbance at 254 nm. (E) Cells of the indicated genotypes and [*psi*⁻]/[*PSI*⁺] states were fivefold serially diluted and spotted onto the indicated plates. (F) The dual-luciferase reporter shown in the schematic was genomically integrated into the strains analyzed in (E). For each strain, eight independent integrants were isolated. Firefly luciferase activity was normalized to Renilla luciferase activity. Fold increases relative to the [*psi*⁻] control are indicated above each bar. *P* values were calculated using an unpaired, two-tailed Student’s t-test (*n* = 8 independent integrants). Black dot, individual data point; error bars, SEM. (G) Cells of the indicated genotypes and [*psi*⁻]/[*PSI*⁺] states were fivefold serially diluted and spotted onto the indicated plates. (H) The dual-luciferase reporter depicted in (F) was genomically integrated into the *upf1*Δ strains analyzed in (G). For each strain, four independent integrants were isolated. Luciferase activity was analyzed as in (F). *P* values were calculated using an unpaired, two-tailed Student’s t-test (*n* = 4 independent integrants). Black dot, individual data point; error bars, SEM.

We first compared basal ISR activity in isogenic [*psi*⁻] and [*PSI*^+^] cells, which differ only in the aggregation state of Sup35. Because ribosome collisions can activate the ISR, we reasoned that collisions at stop codons, if present, would increase ISR activity in [*PSI*^+^] cells. Indeed, [*PSI*^+^] cells activated the ISR in a Gcn1-Gcn20 complex-dependent manner (Fig. 3B), although low levels of eIF2α phosphorylation persisted in [*PSI*^+^] *gcn20*Δ cells. We next restored translation termination fidelity in [*PSI*^+^] cells by expressing an additional, non-aggregating Sup35 variant lacking the N-terminal prion domain (Sup35ΔPrD). These [*PSI*^+^] cells became adenine auxotrophs (Fig. 3A, third row of the spotting assay), indicating that termination at the *ade1-14* premature stop codon was restored. Although Sup35ΔPrD expression conferred a [*psi*⁻]-like adenine-auxotrophic phenotype, endogenous prion-domain-containing Sup35 remained aggregated (Fig. EV3B, third row). Notably, Sup35ΔPrD expression not only restored translation termination fidelity but also reduced the ISR in [*PSI*^+^] cells (Fig. 3C). Thus, ISR activation in [*PSI*^+^] cells results from compromised translation termination rather than from Sup35 aggregation itself. To examine whether compromised translation termination in [*PSI*^+^] cells causes ribosome collision, we compared polysome profiles of [*psi*⁻] and [*PSI*^+^] cells. Because [*PSI*^+^] cells displayed a higher basal level of the ISR (Fig. 3B), we first deleted *GCN2* in both [*psi*⁻] and [*PSI*^+^] strains to eliminate potential effects of differential eIF2α phosphorylation on global translation. [*psi*⁻] *gcn2*Δ and [*PSI*^+^] *gcn2*Δ cells displayed comparable polysome profiles (Fig. EV3C). To reveal collided ribosomes hidden within bulk polysomes, we treated cell lysates with ribonuclease P1 (Ferguson *et al*, 2023). This digestion converts most polysomes into monosomes, whereas collided ribosomes remain as disomes because the mRNA segment between adjacent ribosomes is protected from ribonuclease digestion. Indeed, ribonuclease P1 treatment revealed higher disome levels in [*PSI*^+^] cells (Fig. 3D). Furthermore, Sup35ΔPrD expression markedly reduced these disome levels (Fig. 3D), confirming that the collisions in [*PSI*^+^] cells arise from compromised translation termination. These collided ribosomes likely activate the ISR in [*PSI*^+^] cells through the Gcn1-Gcn20 complex and Gcn2 kinase (Fig. 3B).

### The Gcn1-Gcn20 complex preserves stop codon readthrough in [*PSI*^+^] cells by inhibiting Hel2-dependent RQC

We next examined adenine prototrophy in [*PSI*^+^] cells with impaired ISR. Strikingly, deletion of *GCN1* or *GCN20* produced a [*psi*⁻]-like phenotype (Fig. 3E), despite retention of the aggregated state of Sup35 (Fig. EV3D). In contrast, deletion of the ISR kinase gene *GCN2* did not compromise adenine prototrophy in [*PSI*^+^] cells (Fig. 3E). Thus, the [*psi*⁻]-like phenotype of [*PSI*^+^] *gcn1*Δ and [*PSI*^+^] *gcn20*Δ cells was not caused by loss of the ISR. The compromised adenine prototrophy indicated reduced translational readthrough of the *ade1-14* nonsense mutation. To quantify stop codon readthrough, we used a dual-luciferase reporter containing an internal stop codon between Renilla and firefly luciferasev (Keeling *et al*, 2004). As expected, [*PSI*^+^] cells exhibited higher stop codon readthrough than [*psi*⁻] cells (Fig. 3F). Notably, [*PSI*^+^] *gcn1*Δ and [*PSI*^+^] *gcn20*Δ cells showed reduced stop codon readthrough (Fig. 3F), consistent with their reduced production of functional Ade1 from the *ade1-14* allele (Fig. 3E). To determine why stop codon readthrough was reduced, we first measured frameshifting during translation of the dual-luciferase reporter. Ribosome stalling and collisions can induce frameshifting (Simms *et al*, 2019; Wang *et al*, 2018); if such frameshifting occurred at the internal stop codon of the reporter, it could reduce downstream firefly luciferase expression and thereby lower the apparent readthrough signal. However, deletion of *GCN1* or *GCN20* caused only marginal changes in -1 and +1 frameshifting, which could not account for the reduced stop codon readthrough observed with the 0-frame reporter (Fig. EV3E).

We then reasoned that, when the Gcn1-Gcn20 complex is impaired in [*PSI*^+^] cells, Hel2-mediated clearance of collided ribosomes may compensate for inefficient canonical translation termination, thereby reducing stop codon readthrough. Indeed, deletion of *HEL2* partially restored adenine prototrophy in [*PSI*^+^] *gcn1*Δ and [*PSI*^+^] *gcn20*Δ cells (Fig. EV3F). When nonsense-mediated mRNA decay was impaired by *UPF1* disruption (Leeds *et al*, 1991), *HEL2* deletion fully restored adenine prototrophy in [*PSI*^+^] *gcn1*Δ and [*PSI*^+^] *gcn20*Δ cells (Fig. 3G), indicating recovery of readthrough at the *ade1-14* premature stop codon. Consistently, dual-luciferase assays showed that *HEL2* deletion restored stop codon readthrough in [*PSI*^+^] *gcn1*Δ and [*PSI*^+^] *gcn20*Δ cells to the level observed in [*PSI*^+^] cells (Fig. 3H). In summary, delayed translation termination caused by Sup35 sequestration in [*PSI*^+^] cells results in ribosome collisions that can potentially be resolved by the Hel2-dependent clearance pathway. By protecting ribosomes from this clearance pathway, the Gcn1-Gcn20 complex increases the likelihood of stop codon readthrough, a widely recognized phenotype of [*PSI*^+^] cells.

### The Gcn1-Gcn20 complex limits RQC during physiological translation

Having established that the Gcn1-Gcn20 complex inhibits Hel2-dependent RQC initiation across multiple defined contexts of ribosome pausing, including coding-region stalls and delayed termination at stop codons, we next asked whether this regulatory function also operates within the normal translation pool. Recent studies have revealed that ribosome collisions are widespread during physiological translation under normal conditions (Arpat *et al*, 2020; Han *et al*, 2020; Meydan & Guydosh, 2020; Zhao *et al*, 2021), likely reflecting local variation in elongation rates. We therefore hypothesized that the Gcn1-Gcn20 complex could function as an RQC regulator that sets the threshold for RQC engagement, thereby preventing unnecessary degradation of normal nascent polypeptides during transient ribosome collisions. To measure global RQC initiation in the total translation pool, we took advantage of a reaction that occurs specifically on the 60S•peptidyl-tRNA complex, a unique intermediate generated during RQC. Hel2-dependent RQC initiation generates 60S subunits carrying a peptidyl-tRNA, which are recognized by Rqc2. Rqc2 then appends C-terminal alanine/threonine tails (CAT tails) to the stalled nascent chains (Kostova *et al*, 2017; Osuna *et al*, 2017; Shen *et al*., 2015; Tesina *et al*, 2023); threonine-rich CAT tails then drive detergent-insoluble aggregation (Chang *et al*, 2024; Choe *et al*, 2016; Defenouillere *et al*, 2016; Yonashiro *et al*, 2016) (Fig. 4A). Sis1, a chaperone that associates with these aggregates, can serve as a marker of endogenous stalled polypeptide aggregation (Chang *et al*., 2024; Choe *et al*., 2016; Defenouillere *et al*., 2016; Yonashiro *et al*., 2016). Whereas WT cells efficiently clear endogenous stalled polypeptides through Ltn1-dependent RQC (Fig. EV1A), *ltn1*Δ cells accumulate these aberrant polypeptides. Their co-aggregation with Sis1 can be detected by immunoblotting after semi-denaturing detergent agarose gel electrophoresis (SDD-AGE) (Kryndushkin *et al*, 2003) (Fig. EV4A; compare lanes 1 and 4). This Sis1 aggregate smear was enhanced by overproduction of the CAT-tailing factor Rqc2 (Fig. EV4A, lane 5) (Chang *et al*., 2024; Sitron *et al*, 2020). Notably, Sis1 aggregation increased progressively with cycloheximide-induced translation stress (Fig. EV4B), suggesting that this assay provides a readout of global RQC initiation. Consistent with the severe Sis1 aggregation observed at 10 ng/ml cycloheximide (Fig. EV4B, lane 3), *ltn1*Δ cells overexpressing *RQC2* were highly sensitive to this concentration of cycloheximide (Fig. 4B, fifth row) (Sitron *et al*., 2020). In contrast, *ltn1*Δ cells without *RQC2* overexpression, as well as WT cells with or without *RQC2* overexpression, remained resistant to 10 ng/ml cycloheximide (Fig. 4B). Consistently, these resistant cells displayed only modest Sis1 aggregation (Fig. 4C, lanes 2 and 6). Together, these data suggest that *ltn1*Δ cells overexpressing *RQC2* provide a sensitive system for detecting elevated RQC initiation through two complementary readouts: Sis1 aggregation and impaired cell growth.

**Figure 4.**
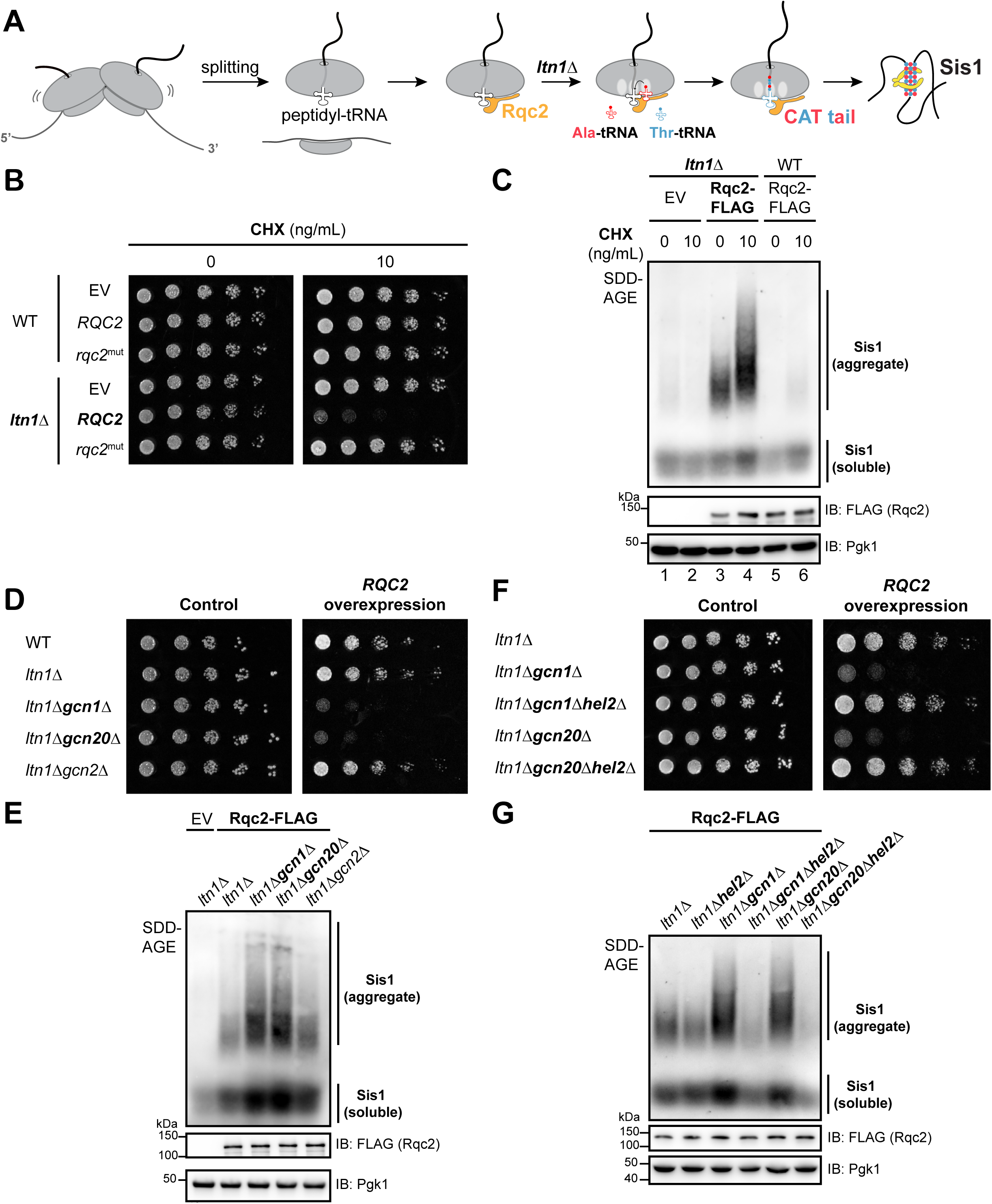
The Gcn1-Gcn20 complex limits RQC during physiological translation (A) Schematic of Rqc2-mediated CAT-tail extension and Sis1-associated aggregation of stalled polypeptides in *ltn1*Δ cells. See also Fig. EV1A, which depicts Ltn1-mediated degradation of stalled polypeptides. (B) Rqc2 or its CAT-tail-deficient mutant, Rqc2^mut^ (D9A, D98A, R99A) (Shen *et al*., 2015), was expressed from the *GAL1* promoter in WT and *ltn1*Δ cells. Cells were fivefold serially diluted and spotted onto galactose plates with or without cycloheximide (CHX). Treatment with 10 ng/ml CHX was generally non-toxic, but caused growth defects when CAT-tail aggregation was enhanced by *LTN1* deletion and *RQC2* overexpression. (C) Rqc2-FLAG was expressed from the *GAL1* promoter in galactose media for 5 h with or without cycloheximide (CHX). Empty vector (EV) served as a control in *ltn1*Δ cells. Cell lysates were analyzed by SDD-AGE (anti-Sis1) and SDS-PAGE (anti-FLAG). Pgk1 served as a loading control. (D) Cells transformed with a *GAL1* promoter-driven *RQC2* overexpression plasmid were serially diluted fivefold and spotted onto glucose (control) and galactose plates. (E) The strains used in (D) were incubated in galactose media for 5 h to induce Rqc2-FLAG expression from the *GAL1* promoter. Cell lysates were analyzed by SDD-AGE (anti-Sis1) and SDS-PAGE (anti-FLAG). EV, empty vector. (F) Cells transformed with a *GAL1* promoter-driven *RQC2* overexpression plasmid were serially diluted fivefold and spotted onto glucose (control) and galactose plates. (G) The strains used in (F) were incubated in galactose media for 6 h to induce Rqc2-FLAG expression from the *GAL1* promoter. Cell lysates were analyzed by SDD-AGE (anti-Sis1) and SDS-PAGE (anti-FLAG).

We next used this sensitized background to test whether the Gcn1-Gcn20 complex suppresses RQC during physiological translation. Accordingly, we overexpressed *RQC2* in *ltn1*Δ*gcn1*Δ and *ltn1*Δ*gcn20*Δ cells and asked whether disruption of the Gcn1-Gcn20 complex enhances Sis1 aggregation and compromises cell growth. Indeed, these mutants exhibited synthetic growth defects upon *RQC2* overexpression (Fig. 4D), whereas *ltn1*Δ*gcn2*Δ cells did not, indicating that the observed toxicity was not caused by loss of the ISR. In addition, the corresponding single-deletion mutants, *gcn1*Δ and *gcn20*Δ, in which the E3 ligase Ltn1 remained intact, did not exhibit synthetic growth defects upon *RQC2* overexpression (Fig. EV4C). This confirms that the toxicity observed in Fig. 4D resulted from accumulation of stalled polypeptides that would otherwise be degraded by Ltn1-dependent RQC. Consistently, Sis1 aggregation was noticeably enhanced in both size and intensity in *RQC2*-overexpressing *ltn1*Δ*gcn1*Δ and *ltn1*Δ*gcn20*Δ cells (Fig. 4E). However, when the RQC pathway remained intact, deletion of *GCN1* or *GCN20* alone did not induce Sis1 aggregation (Fig. EV4D), confirming that the detergent-resistant Sis1 aggregates arose from accumulation of stalled polypeptides. Because CAT tail aggregates are known to induce the heat shock response (HSR) (Brandman *et al*., 2012; Chang *et al*., 2024; Sitron *et al*., 2020), we also examined Hsp104, whose expression is highly responsive to HSR activation (Sanchez & Lindquist, 1990). Indeed, Hsp104 levels were elevated in *ltn1*Δ*gcn1*Δ and *ltn1*Δ*gcn20*Δ cells (Fig. EV4D). Importantly, the *RQC2* overexpression experiments were conducted without translation stressors such as cycloheximide, indicating that physiological translation generates latent RQC targets whose degradative processing is normally suppressed by the Gcn1-Gcn20 complex. When this suppression was lost, these targets entered the Hel2-dependent RQC pathway, as *HEL2* deletion fully restored growth of *ltn1*Δ*gcn1*Δ and *ltn1*Δ*gcn20*Δ cells (Fig. 4F). Consistently, *HEL2* deletion reduced Sis1 aggregation to background levels (Fig. 4G).

## Discussion

Cells maintain proteostasis by balancing protein synthesis, folding, and degradation (Balch *et al*, 2008; Jayaraj *et al*, 2020). This balance deteriorates with age, allowing misfolded proteins to accumulate and form aggregates that contribute to many age-related disorders, particularly neurodegenerative diseases (Hipp *et al*, 2019). Although the mechanisms underlying age-associated proteostasis decline remain incompletely understood, recent studies suggest that ribosome stalling increases with age (Di Fraia *et al*, 2025; Stein *et al*, 2022). If the increased burden of stalled ribosomes exceeds RQC capacity, incomplete nascent polypeptides may escape degradation and cause proteotoxic stress. Indeed, accumulation of stalled polypeptides has been linked to neuromuscular disease (Chu *et al*, 2009; Martin *et al*, 2020), underscoring the importance of efficient RQC in protecting the cellular proteome. A critical challenge in RQC is how to identify rare stalled ribosomes among the vast population of actively translating ribosomes. Cells address this challenge not by detecting stalled ribosomes per se, but by recognizing collided disomes, unique ribosomal complexes formed when trailing ribosomes encounter stalled leading ribosomes (Ikeuchi *et al*., 2019; Juszkiewicz *et al*., 2018). This collision-based recognition mechanism enables highly sensitive RQC. However, recent ribosome profiling studies have identified widespread ribosome collisions during physiological translation (Arpat *et al*., 2020; Di Fraia *et al*., 2025; Han *et al*., 2020; Meydan & Guydosh, 2020; Muller *et al*., 2023; Zhao *et al*., 2021), raising the question of how this sensitive quality-control pathway distinguishes persistent stalls from naturally occurring, potentially transient collisions that may not require RQC engagement.

One possible mechanism for preventing unnecessary RQC is to limit the cellular abundance of the collision sensor Hel2/ZNF598 (Goldman *et al*, 2021). When collision-sensor abundance is low, transiently paused ribosomes may resume elongation before they are recognized and committed to RQC. Consistent with this idea, recent studies showed that Hel2/ZNF598 availability limits RQC efficiency (De La Cruz *et al*, 2025). Here, we reveal an additional layer of RQC control, in which the Gcn1-Gcn20 complex suppresses Hel2-dependent RQC initiation. RQC of the short (AAA)_8_ reporter remains low because the Gcn1-Gcn20 complex opposes RQC initiation (Fig. 2B), whereas the longer (AAA)_10_ reporter undergoes efficient RQC even in the presence of this complex (Fig. 2A). This difference may reflect the ability of Gcn1-Gcn20 to protect transiently collided ribosomes, whereas persistent stalls eventually enter the Hel2-dependent RQC pathway (Fig. 5). Mechanistically, Hel2 may act when Gcn1-Gcn20 dissociates from persistently collided disomes. Alternatively, persistent stalling may generate longer ribosome queues (Goldman *et al*., 2021; Juszkiewicz *et al*., 2018; Matsuo *et al*., 2020), such as trisomes and tetrasomes, whereas transient pausing may preferentially form shorter collided species, such as disomes. If Gcn1-Gcn20 does not fully protect the longer queues, Hel2-dependent ubiquitylation may preferentially occur at persistent stalls rather than at transient pauses.

**Figure 5.**
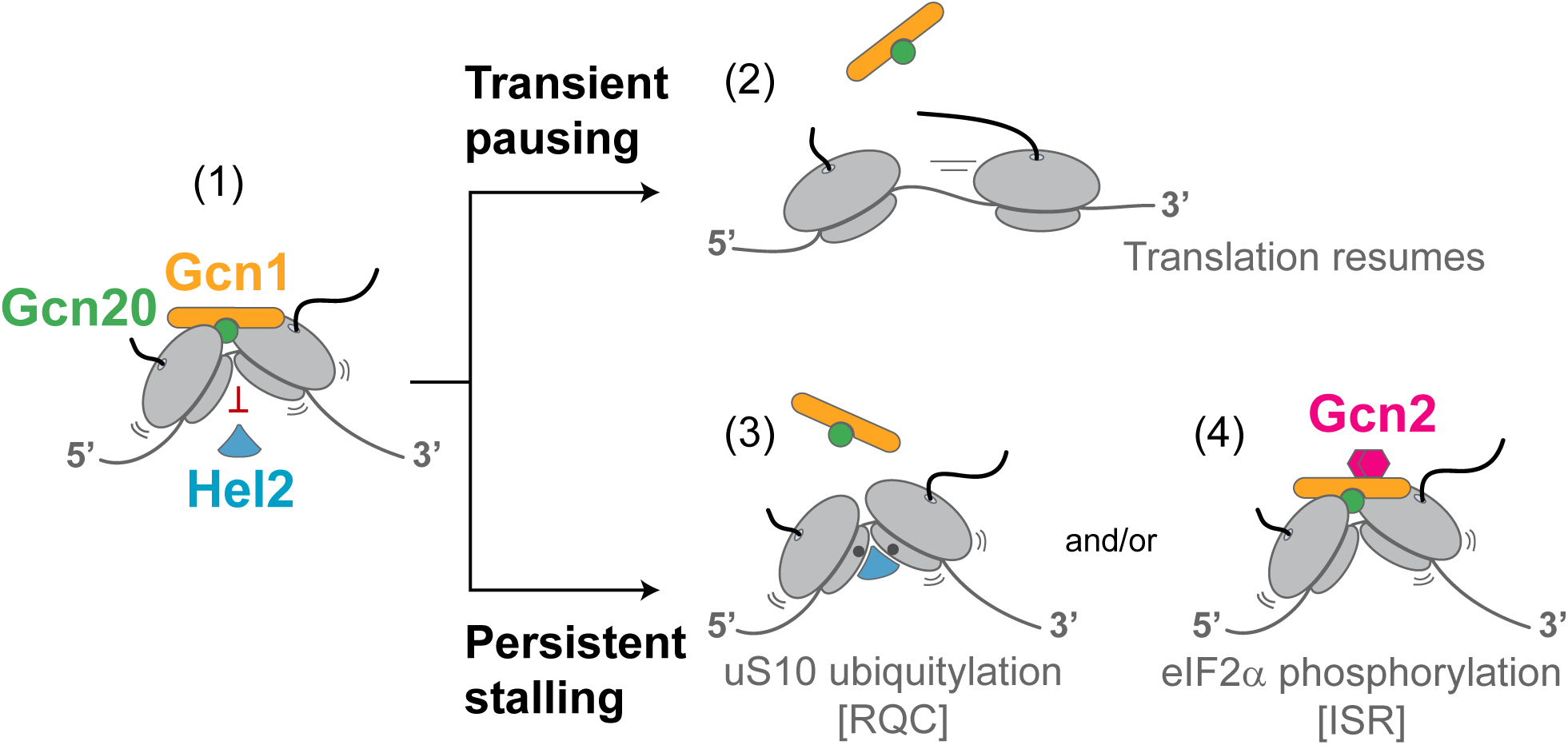
A dual role for the Gcn1-Gcn20 complex: inhibition of RQC and activation of the ISR (1) The Gcn1-Gcn20 complex binds collided ribosomes and inhibits Hel2-mediated RQC initiation. (2) If the leading ribosome is transiently paused, it can resume translation. (3) If the leading ribosome is persistently stalled, it may eventually be processed by Hel2. (4) The Gcn1-Gcn20 complex bound to specific disomes, such as those induced by amino acid starvation, can also activate the ISR kinase Gcn2.

Although antagonistic cross-talk between the ISR and RQC has been reported previously (Yan & Zaher, 2021), the ISR-independent role of the Gcn1-Gcn20 complex in inhibiting RQC had not been recognized, probably because this function is most evident during transient ribosome collisions. In this study, we systematically compared *gcn1*Δ, *gcn20*Δ, and *gcn2*Δ cells under various controlled conditions. The apparent effect of Gcn1-Gcn20 may also depend on the nature of the stalling trigger. During mRNA-damaging stresses, such as RNA alkylation, reactive oxygen species, or UV irradiation (Wu *et al*., 2020; Yan *et al*, 2019), stress severity may determine how frequently ribosomes stall, whereas individual lesions may cause irreversible ribosome arrest. Such persistent stalls may eventually enter the Hel2-dependent RQC pathway regardless of Gcn1-Gcn20-mediated protection. In contrast, the pausing events induced under the conditions used in this study, such as partial amino acid limitation or reduced eRF3 availability, are expected to be more transient and therefore more sensitive to regulation by Gcn1-Gcn20. Under modest translation stress, the Gcn1-Gcn20 complex may serve as a primary defense mechanism that limits premature Hel2-mediated ribosome splitting. As stress becomes more severe, ISR activation would reduce ribosome loading onto mRNAs, thereby limiting further ribosome collisions (Yan & Zaher, 2021).

We also provide evidence that the Gcn1-Gcn20 complex protects the physiological translation pool under normal conditions (Fig. 4). This is consistent with previous studies (Pochopien *et al*., 2021), which isolated a Gcn1-bound disome complex under non-stress conditions (Fig. EV1B). Notably, that complex was isolated by Gcn20 affinity purification, although Gcn20 itself was not structurally resolved. In the resolved disome structure, the leading stalled ribosome contained peptidyl-tRNA in the A-site, uncharged tRNA in the P-site, and the elongation factor eIF5A in the E-site, suggesting that the ribosome may have paused because of impaired elongation at the peptidyl transferase center. Thus, the Gcn1-Gcn20 complex may recognize a broad spectrum of disomes, including those generated by impaired peptide elongation or by amino acid starvation, in which the A-site may be either empty or occupied by an uncharged tRNA. Although the Gcn1-Gcn20 complex can interact with disomes under normal conditions (Pochopien *et al*., 2021), this interaction does not necessarily activate the ISR. First, Gcn2 is present at less than one-fifteenth the abundance of Gcn1 and Gcn20 (Huang *et al*, 2023). Second, Gcn2 recruitment to Gcn1-Gcn20-bound disomes may be regulated, as recent studies showed that Gcn2 is normally bound to non-translating free 60S subunits (Paternoga *et al*., 2025). Notably, collided disomes isolated under normal conditions also contained the Rbg2-Gir2 complex in addition to Gcn1-Gcn20 (Pochopien *et al*., 2021), which may further prevent Gcn2 activation (Wout *et al*, 2009).

Taken together, our study reveals that the Gcn1-Gcn20 complex fine-tunes RQC by preventing its premature initiation, thereby avoiding unnecessary degradation of nascent polypeptides. By sparing transiently collided ribosomes from RQC, this mechanism may allow translation to resume and yield complete protein products.

## Methods

### Yeast culture and transformation

*Saccharomyces cerevisiae* strain BY4741 (MATa *his3*Δ1 *leu2*Δ0 *met15*Δ0 *ura3*Δ0) was used in this study, while strain 74-D694 (MATa *ade1-14 leu2-3,112 his3*Δ200 *trp1-289 ura3-52* [*PIN*^+^]) was used for yeast prion [*PSI*^+^] experiments (Figs. 3 and EV3). All other strains used were derived from BY4741 or 74-D694 and are listed in **Appendix** **Table 1**. Yeast cultures were grown at 30°C unless otherwise specified, in YPD (1% Bacto-yeast extract, 2% Bacto-peptone, 2% glucose), ¼ YPD (0.25% Bacto-yeast extract, 2% Bacto-peptone, 2% glucose), or synthetic drop-out (SD) media consisting of 0.67% yeast nitrogen base supplemented with an appropriate carbon source of either 2% glucose (Sigma, G8270), 2% raffinose (Nacalai Tesque Japan, 30001-15) or 2% galactose (Sigma, G0625). Additional amino acids were supplemented to SD media as follows: 0.04 g/L adenine sulfate (Sigma, A8626), 0.02 g/L L-arginine hydrochloride (Sigma, A5006), 0.1 g/L L-asparagine (Sigma, A8381), 0.1 g/L L-glutamic acid (Sigma, G1626), 0.03 g/L L-lysine hydrochloride (Sigma, L5626), 0.02 g/L L-methionine (Sigma, M9625), 0.05 g/L L-phenylalanine (Sigma, P2126), 0.375 g/L L-serine (Sigma, S4500), 0.2 g/L L-threonine (Sigma, T8625), 0.04 g/L L-tryptophan (Sigma, T0254), 0.03 g/L L-tyrosine (Sigma, T3754), 0.15 g/L L-valine (Sigma, V0500), 0.02 g/L L-histidine hydrochloride (Sigma, 53370), 0.06 g/L leucine (Sigma, L8000), 0.02 g/L uracil (Sigma, U0750). Where indicated, cycloheximide (Sigma, 1810) and copper sulfate (Sigma, 939315) were added, and leucine concentrations were adjusted.

Transformation of plasmids, deletion cassettes, and epitope-tagging cassettes to the yeast genome was performed using the lithium acetate protocol. Briefly, yeast cells in the logarithmic phase were washed twice with water and subjected to heat shock at 42°C for 20 min in a mixture containing polyethylene glycol (PEG3350; Sigma, 202444), lithium acetate (LiAc; Sigma, 62393), single-stranded carrier DNA (ssDNA; Sigma, D1626), and exogenous DNA. Transformants were selected on antibiotic-containing media or on SD media lacking the appropriate amino acid or uracil.

To induce [*PSI*^+^] prion formation in 74-D694 strains, [*psi*⁻] cells were transformed with a plasmid expressing SUP35PrD-RNQ1 under the truncated *SUP35* promoter (Choe *et al*, 2009). White colonies on YPD plates were selected as [*PSI*^+^] candidates and subsequently streaked onto SD plates supplemented with 1 mg/mL 5-fluoroorotic acid (5-FOA; Thermo Fisher Scientific, R0811) to isolate cells that had lost the SUP35PrD-RNQ1 plasmid. The [*PSI*^+^] state was further verified by Sup35PrD-GFP aggregation and by curing on YPD plates containing 1 mM guanidinium chloride.

**Appendix Table 1.** List of yeast strains used in this study.

| Strain | Genotype | Source |
| --- | --- | --- |
| WT (BY4741) | BY4741 (MATa *his3Δ1 leu2Δ0 met15Δ0 ura3Δ0*) | Euroscarf |
| *ltn1Δ* | BY4741 *ltn1Δ::natNT2* | This study |
| *ltn1Δgcn1Δ* | BY4741 *ltn1Δ::natNT2 gcn1Δ::KanMX* | This study |
| *ltn1Δgcn20Δ* | BY4741 *ltn1Δ::natNT2 gcn20Δ::KanMX* | This study |
| *ltn1Δgcn2Δ* | BY4741 *ltn1Δ::natNT2 gcn2Δ::KanMX* | This study |
| *ltn1Δgcn1Δhel2Δ* | BY4741 *ltn1Δ::natNT2 gcn1Δ::KanMX hel2Δ::KILEU2* | This study |
| *ltn1Δgcn20Δhel2Δ* | BY4741 *ltn1Δ::natNT2 gcn20Δ::KanMX hel2Δ::KILEU2* | This study |
| *gcn1Δ* | BY4741 *gcn1Δ::KanMX* | This study |
| *gcn20Δ* | BY4741 *gcn20Δ::KanMX* | This study |
| *hel2Δ* | BY4741 *hel2Δ::KanMX* | This study |
| *mag2Δ* | BY4741 *mag2Δ::KanMX* | This study |
| WT uS3-3HA | BY4741 uS3-3HA-caURA3 | This study |
| *hel2Δ* uS3-3HA | BY4741 *hel2Δ* uS3-3HA-caURA3 | This study |
| *mag2Δ* uS3-3HA | BY4741 *mag2Δ* uS3-3HA-caURA3 | This study |
| WT uS10-3HA | BY4741 uS10-3HA-caURA3 | This study |
| WT uS10-3HA *sui2-S52A* | BY4741 uS10-3HA-caURA3 *sui2-S52A* | This study |
| WT SUI2-2FLAG | BY4741 SUI2-2FLAG-skHIS3 | This study |
| *gcn1Δ* uS10-3HA | BY4741 *gcn1Δ::KanMX* uS10-3HA-caURA3 | This study |
| *gcn20Δ* uS10-3HA | BY4741 *gcn20Δ::KanMX* uS10-3HA-caURA3 | This study |
| *gcn2Δ* uS10-3HA | BY4741 *gcn2Δ::KanMX* uS10-3HA-caURA3 | This study |
| *gcn1Δhel2Δ* uS10-3HA | BY4741 *gcn1Δ::KanMX hel2Δ::NatNT2* uS10-3HA-caURA3 | This study |
| *hel2Δ* uS10-3HA | BY4741 *hel2Δ::NatNT2* uS10-3HA-caURA3 | This study |
| WT HEL2-2FLAG | BY4741 HEL2-2FLAG-skHIS3 | This study |
| *gcn1Δ* HEL2-2FLAG | BY4741 *gcn1Δ::KanMX* HEL2-2FLAG-skHIS3 | This study |
| *gcn20Δ* HEL2-2FLAG | BY4741 *gcn20Δ::KanMX* HEL2-2FLAG-skHIS3 | This study |
| *gcn2Δ* HEL2-2FLAG | BY4741 *gcn2Δ::KanMX* HEL2-2FLAG-skHIS3 | This study |
| WT (74-D694) | MATa *ade1-14 leu2-3,112 his3Δ200 trp1-289 ura3-52 [PIN<sup>+</sup>] [psi<sup>-</sup>]* | Chernoff *et al.*, 1995 |
| [*psi*<sup>-</sup>] WT | 74-D694 [*psi*<sup>-</sup>] | Chernoff *et al.*, 1995 |
| [*PSI*<sup>+</sup>] WT | 74-D694 [*PSI*<sup>+</sup>] | This study |
| [*PSI*<sup>+</sup>] *gcn1Δ* | 74-D694 [*PSI*<sup>+</sup>] *gcn1Δ::KanMX* | This study |
| [*PSI*<sup>+</sup>] *gcn20Δ* | 74-D694 [*PSI*<sup>+</sup>] *gcn20Δ::KanMX* | This study |
| [*PSI*<sup>+</sup>] *gcn2Δ* | 74-D694 [*PSI*<sup>+</sup>] *gcn2Δ::KanMX* | This study |
| [*psi*<sup>-</sup>] *hel2Δ* | 74-D694 [*psi*<sup>-</sup>] *hel2Δ::KILEU2* | This study |
| [*PSI*<sup>+</sup>] *hel2Δ* | 74-D694 [*PSI*<sup>+</sup>] *hel2Δ::KILEU2* | This study |
| [*PSI*<sup>+</sup>] *upf1Δ* | 74-D694 [*PSI*<sup>+</sup>] *upf1Δ::NatNT2* | This study |
| [*PSI*<sup>+</sup>] *upf1Δgcn1Δ* | 74-D694 [*PSI*<sup>+</sup>] *upf1Δ::NatNT2 gcn1Δ::KanMX* | This study |
| [*PSI*<sup>+</sup>] *upf1Δgcn1Δhel2Δ* | 74-D694 [*PSI*<sup>+</sup>] *upf1Δ::NatNT2 gcn1Δ::KanMX hel2Δ::KILEU2* | This study |
| [*PSI*<sup>+</sup>] *upf1Δgcn20Δ* | 74-D694 [*PSI*<sup>+</sup>] *upf1Δ::NatNT2 gcn20Δ::KanMX* | This study |
| [PSI<sup>+</sup>] *upf1Δgcn20Δhel2Δ* | 74-D694 [PSI<sup>+</sup>] *upf1Δ::NatNT2 gcn20Δ::KanMX hel2Δ::KILEU2* | This study |
| [PSI<sup>+</sup>] *gcn1Δhel2Δ* | 74-D694 [PSI<sup>+</sup>] *gcn1Δ::KanMX hel2Δ::KILEU2* | This study |
| [PSI<sup>+</sup>] *gcn20Δhel2Δ* | 74-D694 [PSI<sup>+</sup>] *gcn20Δ::KanMX hel2Δ::KILEU2* | This study |

### Plasmid design

All plasmids used in this study are listed in **Appendix** **Table 2**. Plasmids were constructed using standard molecular cloning techniques with appropriate primers and restriction enzymes and verified by Sanger sequencing (Bio Basic Asia Pacific).

The stalling reporters (AAA)_8_, (AAA)_10_, (AAG)_10_, and (CGA)_3_ were generated by inserting the indicated repeats immediately downstream of GFP using PCR primers. Each PCR fragment was cloned into the yeast vector p416GAL1 using the XbaI and BamHI restriction sites. Following this insert, two tandem HA tags (2xHA) were placed between BamHI and HindIII, followed by mCherry between HindIII and ClaI. Dicodon reporters [(CGA)_2_, (AGA)_2_, CGA-CCG, and AGA-CCA] were constructed by inserting the respective dicodons at codon positions 3 and 4 of GFP (Met-Ser-Codon1-Codon2-GFP) using PCR primers.

The dual-luciferase reporter was constructed to include a premature stop codon (TGA) positioned downstream of Renilla luciferase (inserted via XbaI and BamHI) and upstream of Firefly luciferase (inserted via BamHI and ClaI), as illustrated in Fig. 3F. Both luciferase fragments were amplified by PCR and sequentially cloned into the pRS306 plasmid containing the constitutive GPD promoter and the *CYC1* terminator. The plasmid was linearized with EcoRV prior to integration into the *ura3-52* locus of cells derived from the 74-D694 strain.

When indicated, yeast proteins (Hel2, Mag2, Sup35ΔPrD, uS3 and uS10) were expressed from plasmids, with C-terminal tags introduced via primers to enable immunoblotting of the expressed proteins. *HEL2* and *MAG2* inserts were cloned into pRS413/pRS415 vectors under the control of the *GAL1* promoter using BamHI and HindIII restriction sites. For Sup35ΔPrD expression, two fragments were assembled: the upstream *SUP35* promoter region (flanked by SacI and XbaI) and the C-terminal region of Sup35 (amino acids 253-685; flanked by XbaI and XhoI). These fragments were inserted into pRS416 and subsequently subcloned into pRS306. uS3 and uS10 were cloned into the pRS413/pRS415 backbone using XbaI and XhoI, expressed under the *GPD* promoter. Site-directed mutagenesis with complementary primers was used to substitute catalytic lysine residues with arginine (K212 in uS3; K6 and K8 in uS10).

To quantify translation under leucine depletion, the 5’ untranslated region (5’ UTR) of *GCN4* (nucleotides -404 to -1 relative to the start codon) was inserted upstream of GFP. This region contains four upstream open reading frames (uORFs), with the A of each uORF start codon (ATG) located at positions -361, -293, -176, and -151. The PCR fragment, flanked by BglII and XbaI sites, was cloned into a GFP expression vector under the control of the *GAL1* promoter.

C-terminal 2xFLAG or 3xHA tags were introduced into endogenous genes (HEL2-2FLAG, SUI2-2FLAG, uS10-3HA) by transforming PCR-amplified tagging cassettes into yeast cells. Primers containing 40 bp homology arms upstream and downstream of each target gene were used to amplify the tagging modules from pFA6a-2xFLAG-skHIS3 and pFA6a-3xHA-caURA3 templates.

To mutate the phosphorylation site of endogenous eIF2α (encoded by *SUI2*) from serine to alanine (S52A), we used the pop-in/pop-out (chromosomal integration/excision) method. A fragment containing the *SUI2* promoter and coding sequence for amino acids 1-217, followed by a stop codon, was amplified from genomic DNA. The serine codon (TCC) at position 52 was mutated to alanine (GCC) during PCR amplification using mutagenic primers. The resulting truncated *sui2*-S52A fragment was flanked by BamHI and XhoI restriction sites.

The insert was cloned into the pRS306 plasmid backbone, linearized with MscI within the cloned *sui2* fragment, and transformed into yeast cells. Genomically integrated strains were selected on SD plates lacking uracil. To isolate cells containing only the S52A point mutation without additional modifications, cells were plated on SD-glucose plates supplemented with 0.05 g/L uracil and 1 mg/mL 5-fluoroorotic acid. The presence of the S52A mutation, and the absence of other modifications in the *SUI2* genomic locus were confirmed by sequencing.

**Appendix Table 2.**
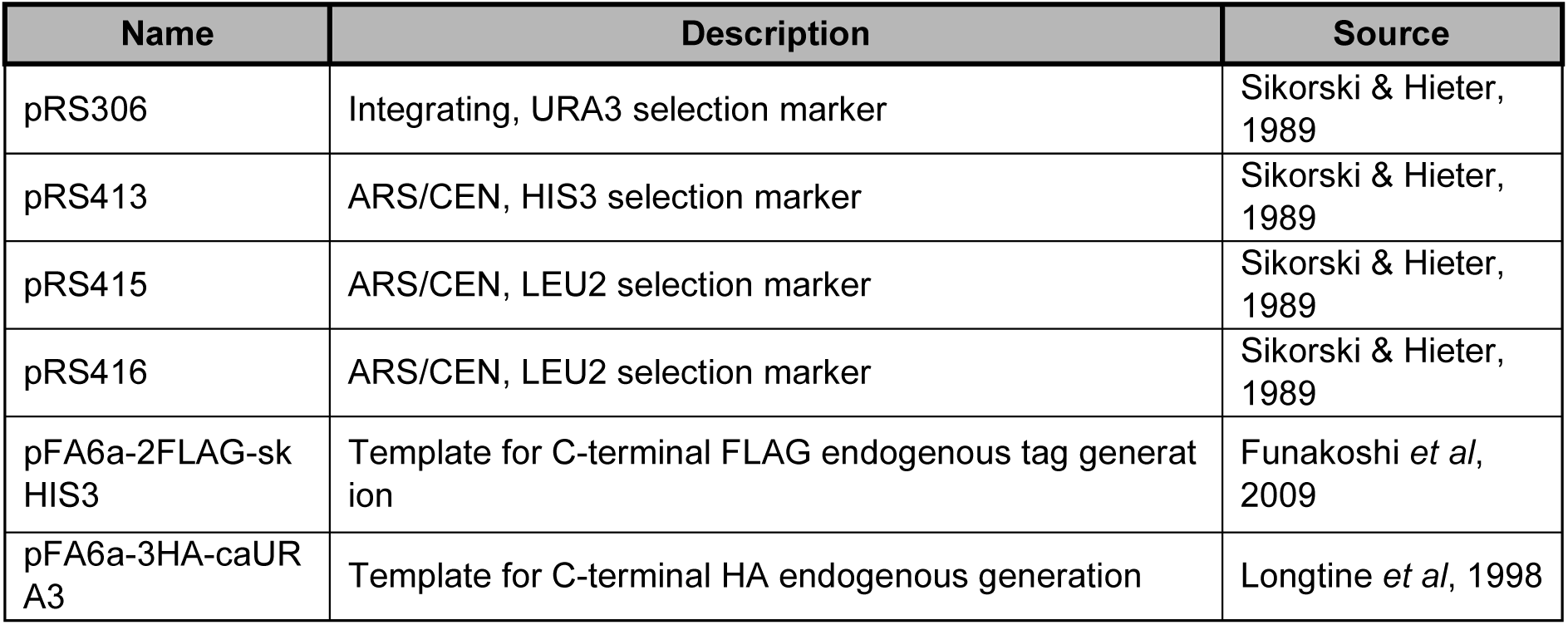
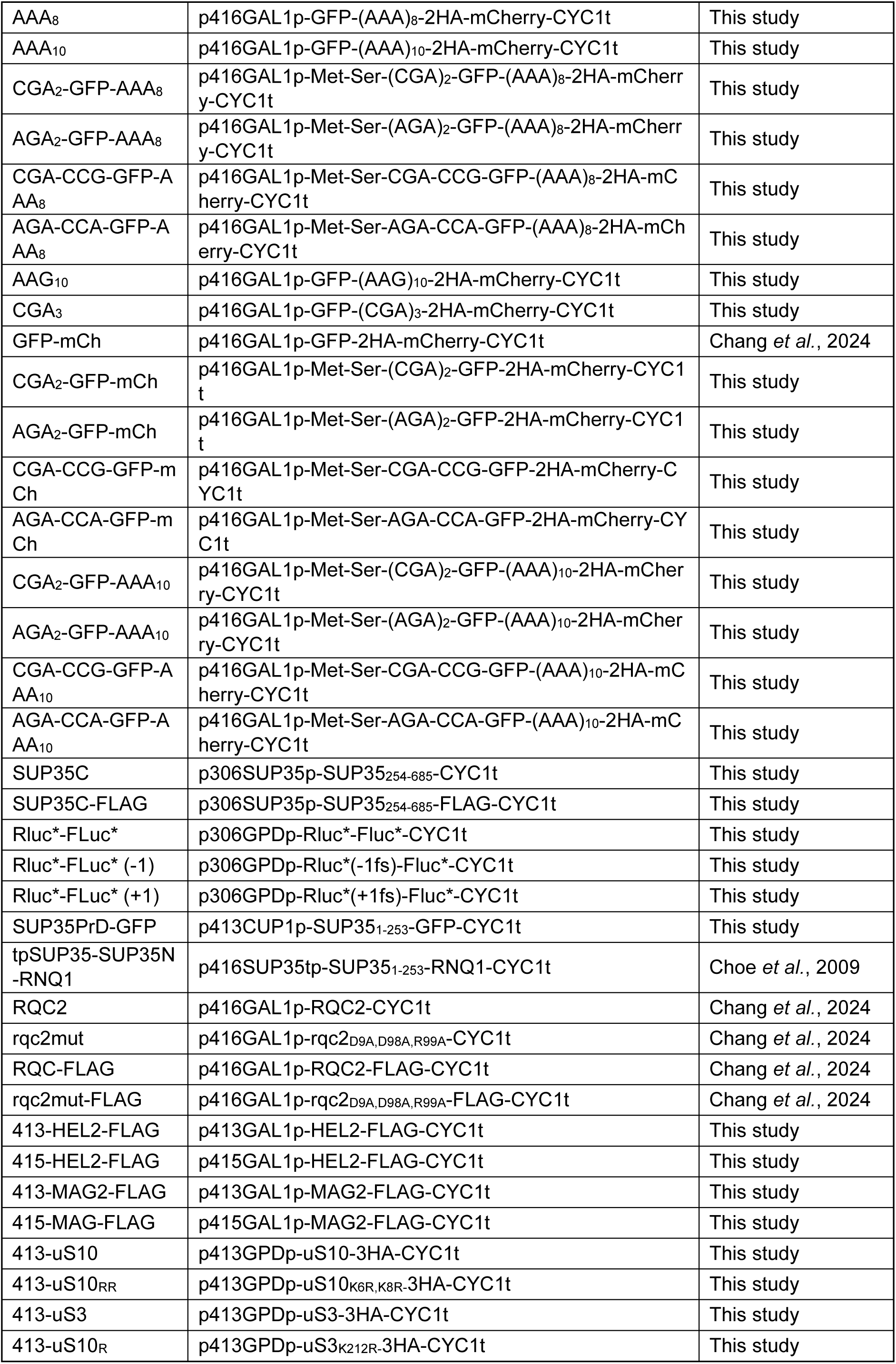
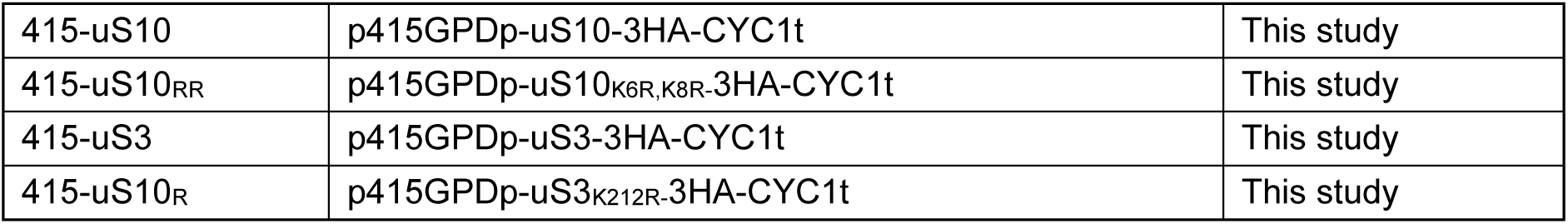
List of plasmids used in this study.

### Yeast lysate preparation for Immunoblotting

Cells were cultured in selective media at 30°C with shaking until reaching an OD_600_ of 0.8–1.0, yielding at least OD_600_ = 50 cell equivalents. For experiments requiring pGAL1 induction, cells were first grown in raffinose-containing SD medium, after which galactose was added to a final concentration of 2%. Where indicated, cycloheximide was added simultaneously with galactose. Unless otherwise specified, all galactose induction experiments were carried out for 5 h.

The liquid culture was chilled on ice and pelleted by centrifugation at 10,000 g for 7 min at 4°C using a Beckman J-25 high-speed centrifuge fitted with a JA-14 rotor (Beckman Coulter). The supernatant was discarded, and the pellet was washed once with ice-cold Milli-Q water, transferred to a pre-chilled 2 mL screw-cap tube, and pelleted again at 10,000 g for 7 min at 4°C. After removing the wash, the final pellet was flash-frozen in liquid nitrogen and stored at -80°C.

To prepare lysates for immunoblotting, 250 μL of acid-washed glass beads (425-600 μm; Sigma, G8772), 400 μL lysis buffer (25 mM Tris-HCl pH 7.4, 150 mM NaCl, 1 mM EDTA, 5% glycerol), and 1x Pierce protease inhibitor (Thermo Scientific, A32965) were added to the frozen yeast pellet. Cells were disrupted at 0°C by glass bead homogenization using a Precellys Evolution homogenizer fitted with a Cryolys adaptor (Bertin Instruments) for four cycles of 40 s, with 90 s pauses between cycles. Cell debris was removed by repeated centrifugation at 2,000 g for 5 min at 4 °C.

Protein concentrations were determined using Bradford reagent (Bio-Rad, 5000006) at 595 nm. Lysates were normalized to equal protein concentrations with additional lysis buffer supplemented with protease inhibitor.

### SDS-PAGE and Immunoblotting

For SDS-PAGE, protein solubilization buffer (PSB; 2% SDS, 5% β-mercaptoethanol, 10% glycerol, 0.25 M Tris-HCl pH 7.4, 0.005% bromophenol blue) was added to lysates, and samples were boiled for 5 min at 98°C. Proteins (20 μg per lane) were separated on 4% stacking/12% resolving Bis-Tris acrylamide gels. Electrophoresis was performed in MES buffer (50 mM MES, 50 mM Tris, 1 mM EDTA, 0.1% SDS, 5 mM sodium bisulfite) at 100 V for 80 min.

Electrotransfer of proteins was performed at 350 mA for 1 h onto 0.45 μm nitrocellulose membranes (Bio-Rad) using Towbin buffer (25 mM Tris, 192 mM glycine, 0.5% SDS, 20% methanol). Membranes were blocked with 3% milk (Sigma, 70166) in Tris-buffered saline containing 0.1% Tween-20 (TBS-T; 20 mM Tris-HCl, 150 mM NaCl, 0.1% w/v Tween-20, pH 7.6) for 1 h, followed by overnight incubation at 4°C with the indicated antibodies at the concentrations specified below in **Appendix** **Table 3**.

**Appendix Table 3.** List of primary and secondary antibodies used in this study.

| Primary antibody | Dilution | Source |
| --- | --- | --- |
| α-GFP | 1:1000 | Roche, 11814460001 |
| α-Pgk1 | 1:2000 | Thermo Fisher Scientific, 459250 |
| α-FLAG | 1:1000 | Sigma-Aldrich, F1804 |
| α-eIF2α / α-Sui2 | 1:2000 | Abcam, Ab32157 |
| α-Sis1 | 1:5000 | Cosmo Bio Japan, COP-080051 |
| α-HA-peroxidase | 1:2000 | Roche, 12013819001 |
| α-Hsp104 | 1:1000 | Abcam, Ab69549 |
| α-mouse IgG | 1:3000 | #7076, Cell Signaling Technology |
| α-rabbit IgG | 1:3000 | #7074, Cell Signaling Technology |

Blots were incubated with the appropriate HRP-conjugated secondary antibodies for 1 h at 25°C. Immunodetection was performed using SuperSignal™ West Pico PLUS (Thermo Scientific, 34580) or SuperSignal™ West Atto Ultimate Sensitivity (Thermo Scientific, A38554) chemiluminescent substrates, and signals were visualized with either ChemiDoc MP Imaging System (Bio-Rad) or Amersham ImageQuant 800 Fluor (Cytiva, 29399484). For reprobing, blots were treated with mild stripping buffer (1.5% w/v glycine, 0.1% SDS, 1% v/v Tween-20, pH 2.2) once for 10 minutes, TBS twice for 10 minutes, and TBS-T twice for 5 minutes. Blots were then blocked and re-incubated with primary antibody as described above. Blots were processed into figures using ImageLab (Bio-Rad) and ImageJ.

### Trichloroacetic Acid (TCA) precipitation

TCA precipitation was used to prepare protein lysates for immunoblotting of post-translational modifications (phosphorylation and ubiquitylation). Yeast cells were grown overnight in SD-glucose medium at 30°C, refreshed to OD_600_ = 0.2 in 10 mL, and cultured to OD_600_ ≍ 0.8. Cells were pelleted at 10,000 g for 2 min at 25°C, and the medium was discarded. The cells were resuspended in SD media of the indicated composition. For amino acid deprivation experiments, leucine concentrations were adjusted as specified in each figure. Where indicated, cycloheximide was added to the medium from a 1 mg/mL stock solution. After 15 min (unless otherwise stated), cells were rapidly chilled in an ice bath to minimize metabolic and biochemical changes during harvesting. Equal numbers of cells (OD_600_) were transferred to pre-chilled tubes. The cells were pelleted at 18,000 g for 6 min at 4°C, the supernatant was discarded, and the pellet was resuspended in 20% TCA then frozen in liquid nitrogen. Samples were stored at -80 °C until lysis.

All subsequent pelleting and washing steps were performed on ice or at 4°C. Samples were thawed completely on ice, pelleted at 10,000 g for 5 min, and the supernatant was removed. Pellets were washed twice with pre-chilled (-20°C) 80% acetone by resuspension and centrifugation at 10,000 g for 5 min each to remove residual TCA. Remaining acetone was allowed to evaporate for 30 min at room temperature. Dried protein pellets were resuspended in alkaline buffer (0.1 M NaOH, 0.1% SDS), followed by addition of an equal volume of TCA protein solubilization buffer (0.5 M Tris-HCl pH 7.4, 4% SDS, 20% glycerol, 0.02% bromophenol blue, 10% β-mercaptoethanol).

To further solubilize and denature proteins, lysates were boiled at 77°C for 7 min. Protein concentrations were determined on a Tecan Infinite M200 Pro plate reader (Tecan, 30050303) using the Pierce™ 660 nm Protein Assay Kit (Thermo Scientific, 22662) with Ionic Detergent Compatibility Reagent (Thermo Scientific, 22663). For each experiment, a standard curve was generated with titrated amounts of bovine serum albumin (BSA; Bio-Rad, 5000201). Sample concentrations were normalized, and 12 μg of total protein was loaded per lane. For eIF2α phosphorylation immunoblotting, membranes were blocked with 3% BSA (Sigma, A2153) in TBS-T.

### Yeast polysome profiling

Yeast cells were grown overnight in YPD, diluted to OD 0.05 in SC media and grown to OD 0.5–0.8, to capture translation during sustained log-phase growth. For Fig. EV1K, cells were washed with pre-warmed leucine-deplete media and starved of leucine for 60 minutes as indicated. Cycloheximide was added to a final concentration of 100 μg/mL for 2 minutes on ice to freeze ribosomes and pelleted at 2,500g for 2 minutes at 4 °C. The pellet was washed once with ice-cold Milli-Q water with 100 μg/mL of Cycloheximide, transferred to a pre-chilled 2 mL screw-cap tube, and pelleted again at 2,500 g for 2 min at 4°C. After removing the wash, the final pellet was flash-frozen in liquid nitrogen, and stored at -80°C or lysed immediately.

To prepare lysates for sucrose gradient analysis, 250 μL of pre-chilled acid-washed glass beads (425-600 mm; Sigma, G8772), 400 μL lysis buffer B (20 mM Tris-HCl pH 7.5, 150 mM NaCl, 5 mM MgCl_2_, and 1 mM DTT), 100 μg/mL Cycloheximide and 1x Pierce protease inhibitor (Thermo Scientific, A32965) were added to the frozen yeast pellet. Cells were disrupted at 0°C by glass bead homogenization using a Precellys Evolution homogenizer fitted with a Cryolys adaptor (Bertin Instruments) for four cycles of 40 s, with 90 s pauses between cycles. The lysates were cleared by centrifugation at 10,000 g for 10 min at 4 °C and normalized to A_260nm_ = 50. Lysates were adjusted to pH 6.5 by adding 7 μL of 300 mM Bis-Tris pH 6.0 per 100 μL of polysome buffered lysate. To digest unprotected mRNA, 10 U/μg of P1 nuclease (M0660S, NEB) was added to each sample and incubated at 30 °C for 60 minutes.

Normalized lysate was layered over a 7–47% sucrose gradient and resolved through ultracentrifugation in a SW41Ti (Beckman) swinging bucket rotor for 3 hours at 35,000 RPM in a 4 °C ultracentrifuge. The resulting polysome gradient was processed by the Biocomp Gradient Master (Model 153) and analyzed at A_254nm_ using the BioRad Econo UV Monitor (BioRad) to measure the polysome profile. Polysome profiles were plotted and compared using GraphPad Prism (version 10.0.3; GraphPad Software).

### Semi-denaturating detergent agarose gel electrophoresis (SDD-AGE)

SDD-AGE was performed to resolve high-molecular-weight aggregates, as previously described with minor modifications. Lysates were mixed 1:1 with 2x SDD-AGE sample buffer (0.1% SDS, 40 mM Tris-acetate, 1 mM EDTA, 10% glycerol, 0.01% bromophenol blue) and incubated at room temperature for 10 min. A total of 30 μg protein was loaded per well onto 1.5% agarose gels containing 0.1% SDS, and electrophoresis was performed in a horizontal tank (Bio-Rad) with SDD-AGE running buffer (0.1% SDS, 40 mM Tris-acetate, 1 mM EDTA) at 75 V for 3 h at 4°C. Proteins were transferred from the gel onto a 0.45 μm nitrocellulose membrane by capillary action using 10 mM Tris-HCl pH 7.4 as the transfer buffer. The system was allowed to stand for a minimum of 12 hours before disassembly, and the membrane washed twice with 10 mM Tris-HCl pH 7.4. Immunoblotting was subsequently performed as described in the SDS-PAGE and immunoblotting section.

### Spotting growth assay

Yeast cells were refreshed in synthetic drop-out (SD) glucose media and diluted to OD_600_ = 0.1. Following 5-fold serial dilutions, 4 μL of each dilution was spotted onto adenine-limited (¼YPD) and adenine-deficient (−Adenine) plates. For galactose induction experiments, transformed yeast cells were refreshed in SD raffinose medium to OD_600_ = 0.1 before serial dilution, and 4 μL of each dilution was spotted onto SD glucose (control/uninduced) and SD galactose (induced) plates with or without 10 ng/mL cycloheximide. Plates were incubated at 30°C for 2-3 days and imaged using ChemiDoc MP Imaging System (Bio-Rad). Images were processed with ImageJ software. For colony color representation in Figures 3 and EV3, ¼YPD spotting images were separated into red, green, and blue (RGB) channels. Using identical region-of-interest (ROI) sizes across channels and spots, average pixel intensities (8-bit) were measured. Values from each ROI were then recombined into an 8-bit RGB format (0,0,0 to 255,255,255) and presented as a colored square.

### Fluorescence microscopy

Live-cell fluorescence imaging was performed using a Nikon Ti Eclipse inverted fluorescence microscope equipped with a Plan-Neofluar 100x/1.30 oil-immersion objective and controlled with NIS-Elements AR software (version 3.22.00; Nikon, Japan). Images were processed using ImageJ software.

### Dual-luciferase assay

Quantitative analysis of stop codon read-through was performed using the Dual-Luciferase® Reporter Assay System (Promega, E1910), which measures Firefly and Renilla luciferase activities. The luciferase reporter was integrated into the *ura3* locus of cells derived from the 74-D694 strain and verified by PCR. For each yeast strain, 4-8 biological replicates were prepared.

Cell lysates and reagents were prepared according to the manufacturer’s instructions. Overnight cultures in exponential phase (OD_600_ = 0.4-0.8) were harvested at room temperature by centrifugation at 10,000 g for 2 min, yielding OD_600_ = 1 cell equivalents. Pelleted cells were washed once with autoclaved Milli-Q water and lysed in 40 μL of 1x passive lysis buffer (PLB) at room temperature. Lysates were kept on ice, and assays were performed at 25°C in 96-well solid white flat-bottom microplates (Corning, CLS3917). Chemiluminescent signals from Firefly and Renilla luciferase were measured sequentially using the Agilent BioTek Cytation 5 Cell Imaging Multimode Reader and normalized to background signals (gain 120, 2 s delay, 1 mm height, 5 s read). The linear range was determined by serial dilution of sample lysates to prevent overflow errors from excessive Renilla luciferase signal, and also ensure that Firefly luciferase signal remained within the detectable range and proportional to lysate abundance. Stop codon read-through was calculated as the ratio of normalized Firefly luciferase activity to normalized Renilla luciferase activity, with Firefly reporting read-through and Renilla reporting total reporter translation. Data points were averaged, and standard error of mean (SEM) was calculated. Graphs were generated using GraphPad Prism (version 10.0.3; GraphPad Software).

### Statistical Analysis

Details of statistical analyses are provided in the figure legends. All analyses were performed using Microsoft Excel and GraphPad Prism (version 10.0.3; GraphPad Software). Statistical significance was assessed using Student’s t-tests (two-tailed, paired/unpaired) with *P* values reported to three significant figures, in scientific notation where applicable. Error bars, where indicated, represent the standard error of mean (SEM).

## Acknowledgments

We thank Que Kong and Wei Ma for their technical guidance. This research is supported by the Ministry of Education, Singapore, under its Academic Research Fund Tier 1 (RT20/23 and RG114/23; Y.-J.C.) and Nanyang Technological University, Singapore, under its Nanyang Assistant Professorship Start-Up Grant (Y.-J.C.).

## Author contributions

Conceptualization, Y.-J.C.; Methodology, T.Y.K.C., W.Y.X.C., M.-J.Y., and Y.-J.C.; Investigation,

T.Y.K.C., W.Y.X.C., M.-J.Y.; Resources, Y.-J.C., Writing – Original Draft, Y.-J.C.; Writing – Review & Editing, T.Y.K.C., W.Y.X.C., and Y.-J.C.; Visualization, T.Y.K.C., W.Y.X.C., and Y.-J.C.; Funding Acquisition, Y.-J.C.; Supervision, Y.-J.C.

---

**Figure EV1.**
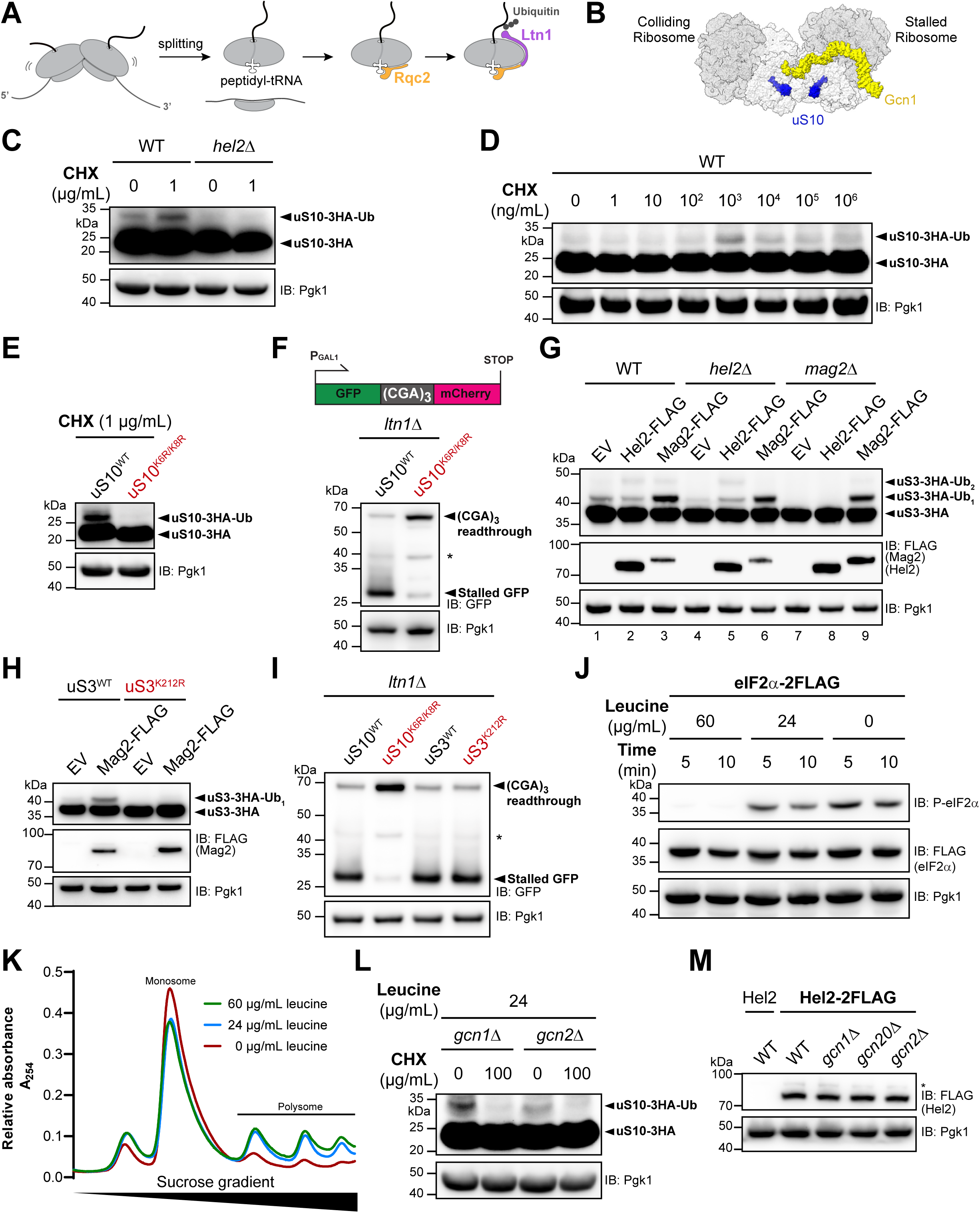
Gcn1 inhibits RQC during partial amino acid depletion (A) Schematic of the ribosome-associated quality control (RQC) pathway. (B) Structure of Gcn1 bound to collided disomes, with uS10 highlighted in blue. The model was rendered from PDB 7NRC and 7NRD (Pochopien *et al*., 2021). (C) Genomic uS10/*RPS20* was C-terminally tagged with a 3×HA epitope in the WT strain, and *HEL2* was deleted in this background. Cells were incubated for 15 min in media with or without cycloheximide (CHX), and uS10 ubiquitylation was assessed by anti-HA immunoblotting. Pgk1 served as a loading control. IB, immunoblotting; Ub, ubiquitin. (D) WT uS10-3HA cells described in (C) were precultured in nutrient-rich synthetic complete medium, transferred to the same media containing increasing concentrations of cycloheximide (CHX; 10-fold increments), and incubated for 15 min. uS10 ubiquitylation was assessed by anti-HA immunoblotting. (E) C-terminally 3×HA-tagged wild-type uS10 (uS10^WT^) and mutant uS10 carrying K6R and K8R substitutions (uS10^K6R/K8R^) were expressed from plasmids under the GPD promoter in cells lacking genomic uS10. uS10 ubiquitylation was induced by treatment with 1 μg/ml CHX for 15 min. (F) *LTN1* was additionally deleted from the strains described in (E). The resulting strains were transformed with the (CGA)_3_ stalling reporter, which was expressed from the *GAL1* promoter for 5 h. Cell lysates were analyzed by anti-GFP immunoblotting. * degradation product of the readthrough protein. (G) Genomic uS3/*RPS3* was C-terminally tagged with a 3×HA epitope in the WT strain, and *HEL2* or *MAG2* was deleted in this background. Hel2-FLAG or Mag2-FLAG was expressed from the *GAL1* promoter in the indicated uS3-3HA strains. Empty vector (EV) served as a control in each strain. uS3 ubiquitylation was assessed by anti-HA immunoblotting. Hel2-FLAG and Mag2-FLAG expression was confirmed by anti-FLAG immunoblotting. Ub_1_, mono-ubiquitylated uS3; Ub_2_, di-ubiquitylated uS3. (H) Mag2 -FLAG was expressed from the *GAL1* promoter in cells lacking endogenous uS3/*RPS3* and expressing 3×HA-tagged wild-type uS3 (uS3^WT^) or mutant uS3 (uS3^K212R^) from plasmids driven by the GPD promoter. EV, empty vector; Ub_1_, mono-ubiquitylated uS3. (I) *LTN1* was additionally deleted from the strains described in (H). The resulting strains were transformed with the (CGA)_3_ stalling reporter, which was expressed from the *GAL1* promoter for 5 h. Cell lysates were analyzed by anti-GFP immunoblotting. * degradation product of the readthrough protein. (J) Genomic eIF2α/*SUI2* was C-terminally tagged with a 2×FLAG epitope in the WT strain. Cells cultured in synthetic complete media containing 60 μg/ml leucine were collected by centrifugation, resuspended in media containing 60, 24, or 0 μg/ml leucine, and incubated for 5 or 10 min before lysate preparation. ISR activation was assessed by immunoblotting with an antibody against phosphorylated eIF2α (P-eIF2α). Total eIF2α levels were analyzed by anti-FLAG immunoblotting. (K) WT cells cultured in nutrient-rich synthetic complete medium were collected by centrifugation, washed once with medium lacking leucine, and resuspended in media containing 60, 24, or 0 μg/ml leucine. After incubation at 30°C for 1 h, lysates were normalized based on absorbance at 260 nm and layered onto 7–47% sucrose gradients. Polysome profiles were analyzed by absorbance at 254 nm. (L) *gcn1*Δ and *gcn2*Δ cells carrying genomic uS10 tagged with 3×HA were cultured in leucine-rich media (60 μg/ml). After collection by centrifugation, cells were resuspended in leucine-limited media (24 μg/ml) with or without cycloheximide (CHX). After incubation for 15 min, uS10 ubiquitylation was assessed by anti-HA immunoblotting. (M) Genomic *HEL2* was C-terminally tagged with a 2×FLAG epitope in WT, *gcn1*Δ, *gcn20*Δ, and *gcn2*Δ strains. Lysate from WT cells lacking *HEL2*-2×FLAG was included in the first lane as an untagged control. Cells were cultured in nutrient-rich synthetic complete medium at 30°C under normal conditions and collected during logarithmic growth.

**Figure EV2.**
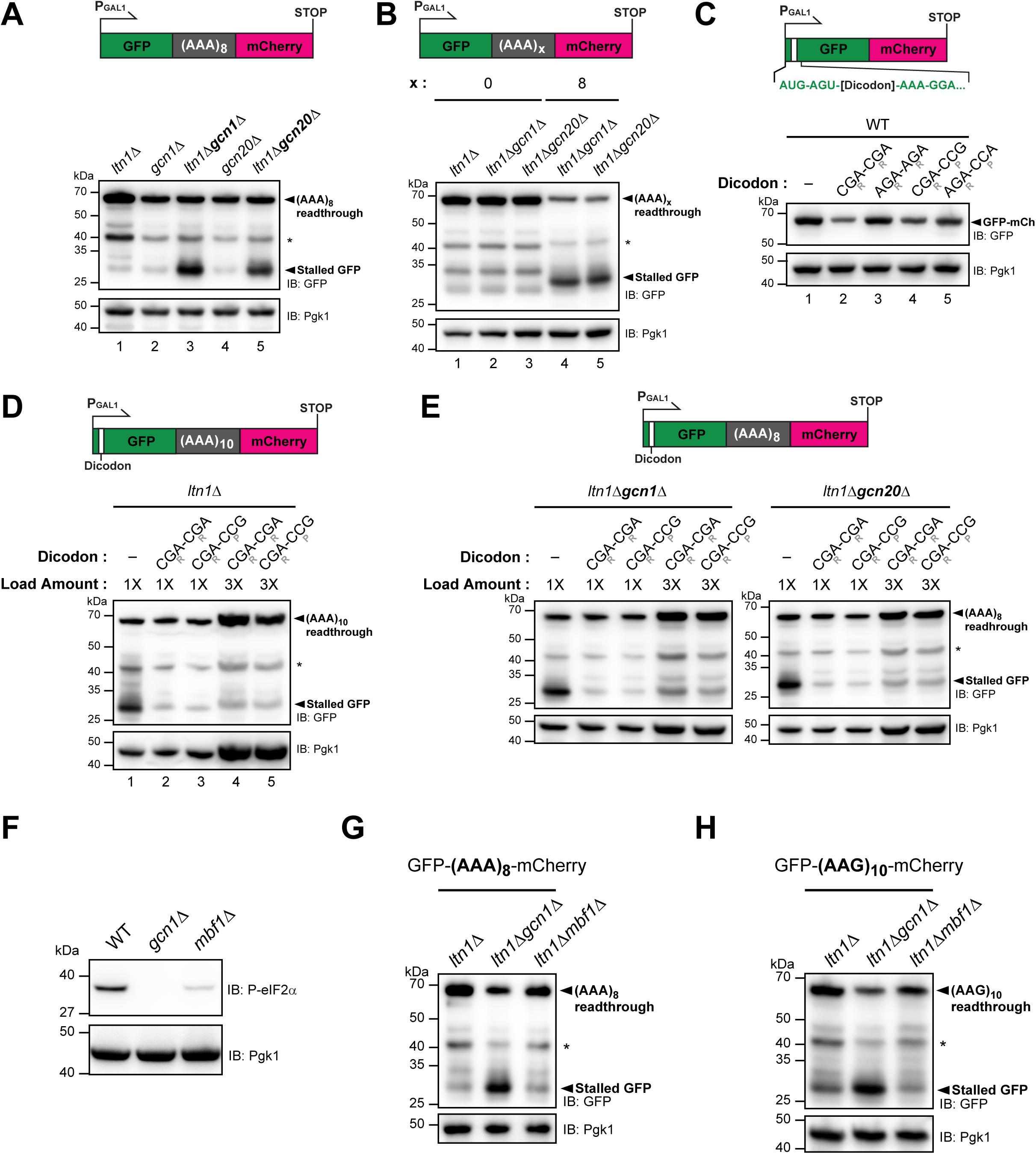
The Gcn1-Gcn20 complex inhibits RQC during translation of polyadenosine tracts (A) The (AAA)_8_ stalling reporter was expressed from the *GAL1* promoter in the indicated strains. Cell lysates were analyzed by anti-GFP immunoblotting. Pgk1 served as a loading control. IB, immunoblotting; * degradation product of the readthrough protein. (B) (AAA)_x_ denotes x repeats of the AAA codon (x = 0 or 8). The reporters were expressed from the *GAL1* promoter in the indicated strains. The (AAA)_8_ reporter was included to indicate the position of stalled GFP. Cell lysates were analyzed by anti-GFP immunoblotting. (C) Translation-inhibitory dicodons (CGA-CGA and CGA-CCG) or their synonymous counterparts (AGA-AGA and AGA-CCA) were inserted at the third codon position of the GFP-mCherry fusion reporter, after the AUG start codon and AGU. CGA and AGA encode arginine (R); CCG and CCA encode proline (P). Reporters were expressed in WT cells. The GFP-mCherry reporter without dicodon insertion was included in the first lane. (D) Lysates from cells expressing the (AAA)_10_ reporters containing the inhibitory dicodons CGA-CGA or CGA-CCG were loaded at threefold higher amounts in lanes 4 and 5 than lysate from cells expressing the reporter without dicodon insertion in lane 1. (E) In both panels, lysates from cells expressing the (AAA)_8_ reporters containing the inhibitory dicodons CGA-CGA or CGA-CCG were loaded at threefold higher amounts in lanes 4 and 5 than lysates from cells expressing the reporter without dicodon insertion in lane 1. All constructs were expressed in *ltn1*Δ*gcn1*Δ cells in the left panel and *ltn1*Δ*gcn20*Δ cells in the right panel. (F) The indicated strains were cultured in nutrient-rich synthetic complete medium under normal conditions. Basal ISR activity was assessed by immunoblotting with an antibody against phosphorylated eIF2α. (G-H) The (AAA)_8_ reporter (G) and the (AAG)_10_ reporter (H) were expressed from the *GAL1* promoter in the indicated strains. RQC activity of each reporter was assessed by anti-GFP immunoblotting.

**Figure EV3.**
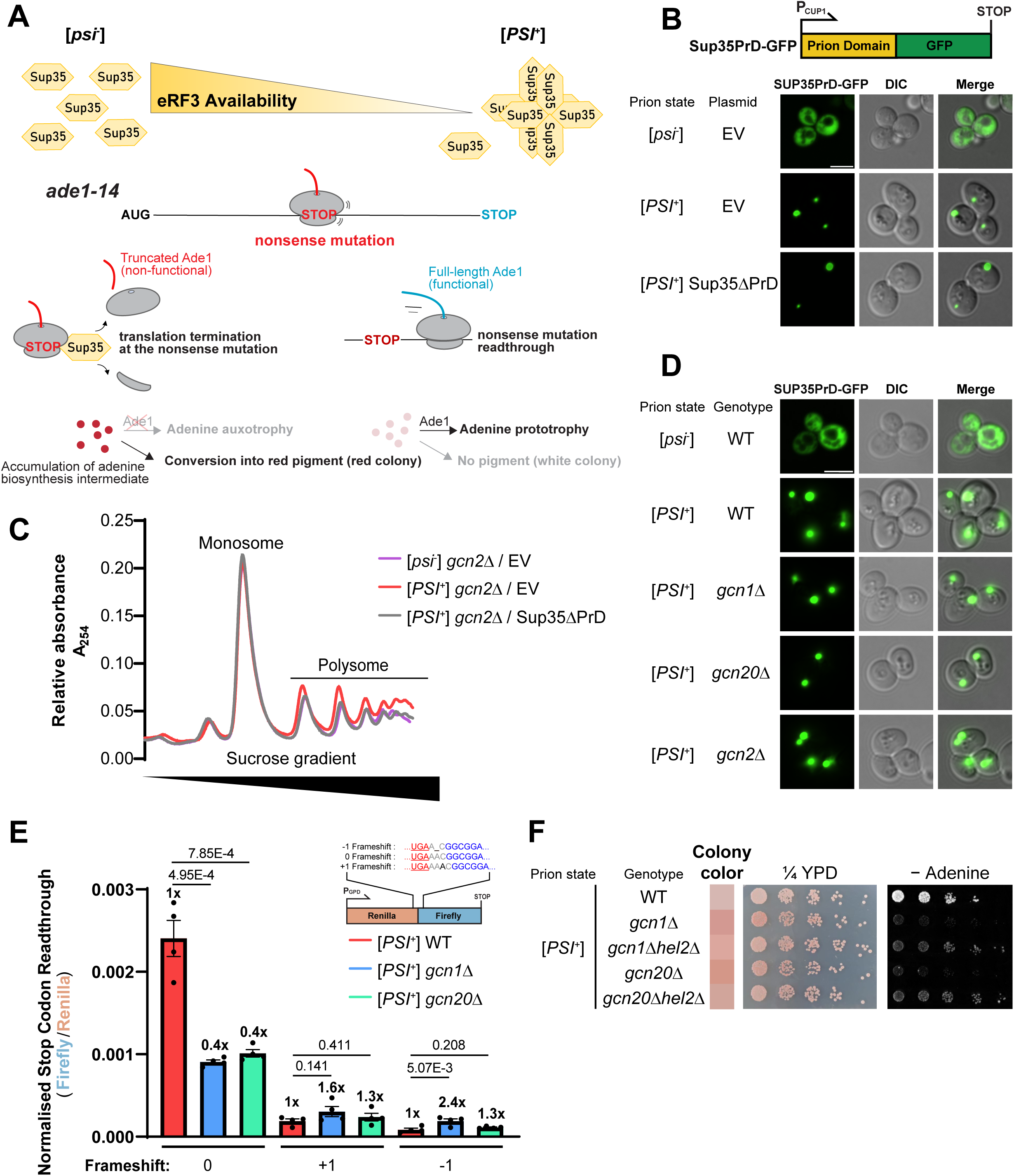
The Gcn1-Gcn20 complex preserves stop codon readthrough in [*PSI*^+^] cells (A) Schematic of the yeast prion [*PSI*^+^] phenotype. [*psi*^-^] cells cannot produce functional Ade1, leading to accumulation of adenine biosynthesis intermediates upstream of the Ade1-catalyzed reaction. These intermediates form a red pigment, resulting in red colonies. In contrast, [*PSI*^+^] cells produce functional Ade1, preventing accumulation of these intermediates and yielding white colonies. (B) Confirmation of the [*PSI*^+^] state by fluorescence microscopy of Sup35PrD-GFP. Sup35PrD-GFP, consisting of the N-terminal prion domain of Sup35 fused to GFP, was expressed from the *CUP1* promoter for 4 h after induction with 100 μM CuSO_4_ in [*psi*^-^] and [*PSI*^+^] cells. In [*PSI*^+^] cells, endogenous Sup35 aggregates template Sup35PrD-GFP aggregation, producing green puncta (Patino *et al*., 1996). In contrast, Sup35PrD-GFP remains diffuse in [*psi*^-^] cells. Although expression of Sup35ΔPrD abolishes the adenine-prototrophy phenotype of [*PSI*^+^] cells (see Fig. 3A), the aggregation state of endogenous Sup35 remains unaffected, as indicated by green puncta in the third row. EV, empty vector; DIC, differential interference contrast microscopy; Scale bar, 5 μm. (C) [*psi*⁻] *gcn2*Δ and [*PSI*⁺] *gcn2*Δ cells were transformed with either empty vector (EV) or a Sup35ΔPrD expression plasmid, as indicated. Cell lysates were resolved through 7-47% sucrose gradients. Polysome profiles were analyzed by absorbance at 254 nm. The same lysates were used for Fig. 3D after P1 nuclease digestion. (D) A Sup35PrD-GFP expression plasmid was transformed into the indicated strains. Sup35PrD-GFP expression from the *CUP1* promoter was induced with 100 μM CuSO_4_ for 4 h. Scale bar, 5 μm. (E) -1 and +1 frameshift dual-luciferase reporters were generated by deleting or inserting one nucleotide after the stop codon UGA between Renilla and firefly luciferase. The original reporter used in Fig. 3F is referred to as the 0-frame reporter. Each reporter was genomically integrated into [*PSI*^+^] WT, [*PSI*^+^] *gcn1*Δ, and [*PSI*^+^] *gcn20*Δ cells. Four independent integrants were isolated for each strain and reporter. Firefly luciferase activity was normalized to Renilla luciferase activity. Fold increases relative to [*PSI*^+^] WT cells are indicated above each bar. *P* values were calculated using an unpaired, two-tailed Student’s t-test (*n* = 4 independent integrants). Black dots, individual data points; error bars, SEM. (F) [*PSI*⁺] cells of the indicated genotypes were fivefold serially diluted and spotted onto adenine-limited (¼YPD) and adenine-deficient (-Adenine) plates.

**Figure EV4.**
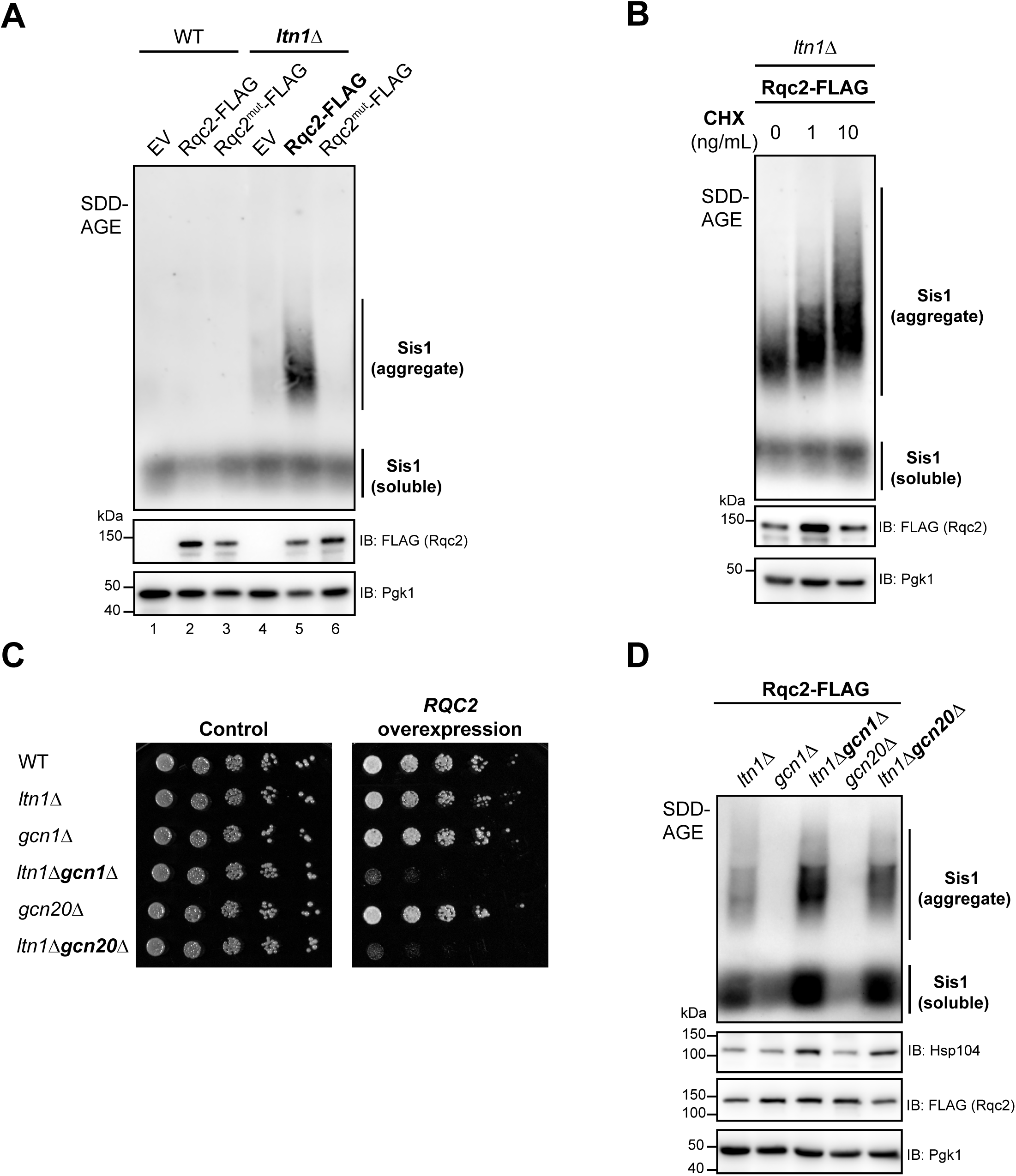
The Gcn1-Gcn20 complex limits RQC during physiological translation (A) Rqc2-FLAG or its CAT-tail-deficient mutant, Rqc2^mut^-FLAG (D9A, D98A, R99A) (Shen *et al*., 2015), was expressed from the *GAL1* promoter in WT and *ltn1*Δ cells for 5 h. Empty vector (EV) served as a control. Cell lysates were analyzed by SDD-AGE using anti-Sis1 immunoblotting and by SDS-PAGE using anti-FLAG immunoblotting. Sis1-associated aggregates appear as a smear on SDD-AGE immunoblots. Pgk1 served as a loading control. (B) Rqc2-FLAG was expressed from the *GAL1* promoter in *ltn1*Δ cells for 5 h in galactose media supplemented with 0, 1 or 10 ng/mL cycloheximide (CHX). Cell lysates were analyzed by SDD-AGE (anti-Sis1) and SDS-PAGE (anti-FLAG). (C) Cells transformed with the plasmid overexpressing *RQC2* from the *GAL1* promoter were serially diluted fivefold and spotted onto glucose (control) and galactose plates. (D) Rqc2-FLAG was expressed from the *GAL1* promoter for 5 h in the indicated cells. Cell lysates were analyzed by SDD-AGE (anti-Sis1) and SDS-PAGE (anti-Hsp104 and anti-FLAG).

